# Obesity phenotypes are preserved in intestinal stem cell enteroids from morbidly obese patients

**DOI:** 10.1101/2020.05.29.123737

**Authors:** Nesrin M. Hasan, Kelli F. Johnson, Jianyi Yin, Nicholas W. Baetz, Vadim Sherman, Sarah E. Blutt, Mary K. Estes, Vivek Kumbhari, Nicholas C. Zachos, Olga Kovbasnjuk

## Abstract

Obesity and obesity-related comorbidities are significant health care challenges. Bariatric surgery (BS) is the most effective therapy for treating obesity and type 2 diabetes. A barrier in the development of therapeutic alternatives is incomplete mechanistic understanding of the benefits of BS and the lack of human intestinal models that recapitulate the pathophysiology of obesity. Using adult intestinal stem cell-derived enteroid cultures established from healthy lean subjects and morbidly obese patients, including post-BS cases, four phenotypes correlating patient BMI and intestinal glucose absorption were identified suggesting that enteroids retain patient phenotype heterogeneity associated with healthy and diseased state. In a sub-population of obese patients, increased dietary glucose absorption and gluconeogenesis was due to significantly higher expression of intestinal carbohydrate transporters (SGLT1, GLUT2 and GLUT5) and gluconeogenic enzymes (PEPCK1 and G6Pase) compared to enteroids from lean subjects that demonstrated low glucose absorption and lacked gluconeogenesis. Enteroids established from successful BS cases exhibited low glucose absorption similar to that observed in lean subjects. These data show that human enteroids preserve the patient phenotype in long-term cultures and represent a reliable preclinical model to study the heterogeneity of the obesity mechanisms, which is necessary to determine the efficacy of therapeutic interventions.

## Introduction

Obesity is a rapidly growing global health problem that is associated with many obesity related comorbidities including insulin resistance, type 2 diabetes (T2D), cardiovascular diseases, and cancer [1, 2]. According to the World Health Organization, ∼ 39% of adults are overweight or obese. The increasing prevalence of obesity is due to a complex mixture of genetic factors and energy imbalance. Dieting and exercise can yield weight loss; however, it is often modest and transient, with limited improvement in glycemic control [3, 4, 5].

Bariatric surgery (BS), which influences intestinal sugar transport and metabolism [6, 7, 8], is an effective option to reverse obesity and T2D. Serum glucose concentrations decrease prior to significant weight loss immediately following BS [9, 10]. Recent studies also show that glucose absorption in the proximal intestine is elevated in morbid obesity, leading to hyperglycemia [11]. However, studies designed to characterize and measure altered glucose absorption in the small intestine from obese patients compared to healthy lean subjects are limited.

Glucose is the main energy source of mammalian cells. Glucose transport is mediated through Na^+^-coupled glucose cotransporters (SGLTs) and facilitative glucose transporters (GLUTs). At low luminal glucose concentration (≤25 mM), glucose is taken up by enterocytes via SGLT1 expressed at the brush border membrane (BBM) [12, 13]. GLUT2 is expressed at the basolateral membrane (BLM) and transports glucose and fructose into the blood [14, 15]. At higher luminal glucose concentrations (≥50 mM), such as following a meal, GLUT2 traffics to the BBM, resulting in uptake of the additional luminal glucose [16]. Fructose, another dietary sugar, enters enterocytes through GLUT5 at the BBM [17]. At high fructose conditions, GLUT2 also localizes to the BBM [18] and contributes to fructose uptake. GLUT1 is expressed at the BLM of enterocytes and has been linked to obesity and diabetes [7, 19], however its role in intestinal epithelial glucose transport is not well understood.

Significantly increased expression of intestinal sugar transporters (SGLT1, GLUT2, and GLUT5) has been observed in different animal models of obesity and/or diabetes [20, 21, 22]. Interestingly, studies in humans indicate heterogeneous expression [23] and cellular distribution of these major sugar transporters in obesity. GLUT2 was significantly upregulated and continuously present at the BBM of enterocytes in a majority (∼72%) of obese individuals in several studies, while it was absent from the BBM in lean subjects and a subset of obese individuals [24, 25]. Conversely, in another study, GLUT2 was not localized apically in duodenal biopsies of overweight T2D subjects [21]. Elevated SGLT1 expression in enterocytes from morbidly obese patients positively correlated with elevated blood glucose; however, the expression of SGLT1 was variable across the studied cohort [11].

*De novo* glucose production by intestinal epithelia via gluconeogenesis might also contribute to the increased serum glucose levels. Gluconeogenesis primarily occurs in the liver [26], hence most studies have focused on hepatic glucose production in normal physiology and obesity [27, 28]. Several studies in animal models have focused on characterizing the role of intestinal gluconeogenesis, however its role in obesity is unclear. When luminal fructose levels are low, the small intestine metabolizes ∼90% of dietary fructose, whereas high doses result in fructose overflow to the liver [29]. Other studies have shown that intestinal gluconeogenesis is involved in gut-brain axis signaling and regulation of energy homeostasis [30].

Overall, these findings are consistent with the concept that increased intestinal glucose absorption and possibly intestinal gluconeogenesis are associated with hyperglycemia in obesity and T2D. Analysis of human samples imply that the expression and localization of the main glucose and/or fructose transporters is heterogeneous in morbid obesity [11, 21, 23, 24, 25]. On account of this, the molecular classification of inter-individual differences may be a key predictor for the outcome of BS or calorie restricted diet. Therefore, mechanistic understanding of the changes in intestinal glucose homeostasis in obesity might be important for generating effective novel therapeutic options and less invasive alternatives to BS.

In order to determine whether the intestinal epithelium functionally differs in regard to glucose absorption and metabolism among lean subjects, overweight/obese patients, and post-BS patients, it is crucial to study relevant *ex vivo* models. We utilized novel intestinal stem cell-derived epithelial enteroid cultures [31, 32, 33, 34] to investigate the contribution of intestinal glucose absorption and gluconeogenesis to the systemic glucose load in lean subjects and in obese patients, including post-BS cases. This is the first study to address the relationship between BMI and the expression and function of sugar transporters and gluconeogenic enzymes in human enteroids and provide evidence that the heterogeneity of obesity-related phenotypes is retained in human enteroids.

## Materials and Methods

Protocols are described in the supplementary methods section.

## Results

### Analysis of the relationship between subject’s BMI and glucose absorption in enteroid cultures demonstrates four phenotypes

Duodenal and/or jejunal small intestinal tissue samples were collected from eighteen subjects (Table S1) and enteroid cultures were generated from twenty-three tissue samples to study the mechanistic differences in dietary sugar absorption and metabolism as a function of BMI. The absorption (defined as transcellular transport of nutrients from the lumen across the intestinal epithelium to the blood) of dietary sugars, mainly occurs in the proximal small intestine (duodenum and jejunum). Enterocytes of small intestinal villi, but not immature enterocytes in the crypts, are mainly responsible for carbohydrate absorption [35]. To measure the sugar absorption, we used confluent enteroid monolayers (EM) grown on Transwell filters which allow access to both apical and basolateral surfaces [32, 33], and modeled as crypt-like or villus-like epithelia using undifferentiated (UD) or differentiated (DF) EM, respectively [31, 32, 36].

DF-EM, derived from 23 tissue samples, were exposed to 25 mM glucose apically and changes in the concentration of glucose were measured in basolateral media as an indicator of glucose absorption. Four phenotypes representing the correlation between BMI and glucose absorption (GA) were found (Fig. 1A and Table S1): BMI^lean^GA^low^, BMI^overweight^GA^high^, BMI^obese^GA^high^ and BMI^obese^GA^low^. Enteroids from obese subjects had two glucose absorption phenotypes (GA^low^ or GA^high^). Enteroids from overweight patients exhibited similar glucose absorption to the BMI^obese^GA^high^ group. These data provide evidence for the heterogeneity of glucose absorption in morbidly obese subjects and suggest that elevated intestinal glucose absorption in a subset of obese subjects may significantly contribute to hyperglycemia.

**Fig. 1.**
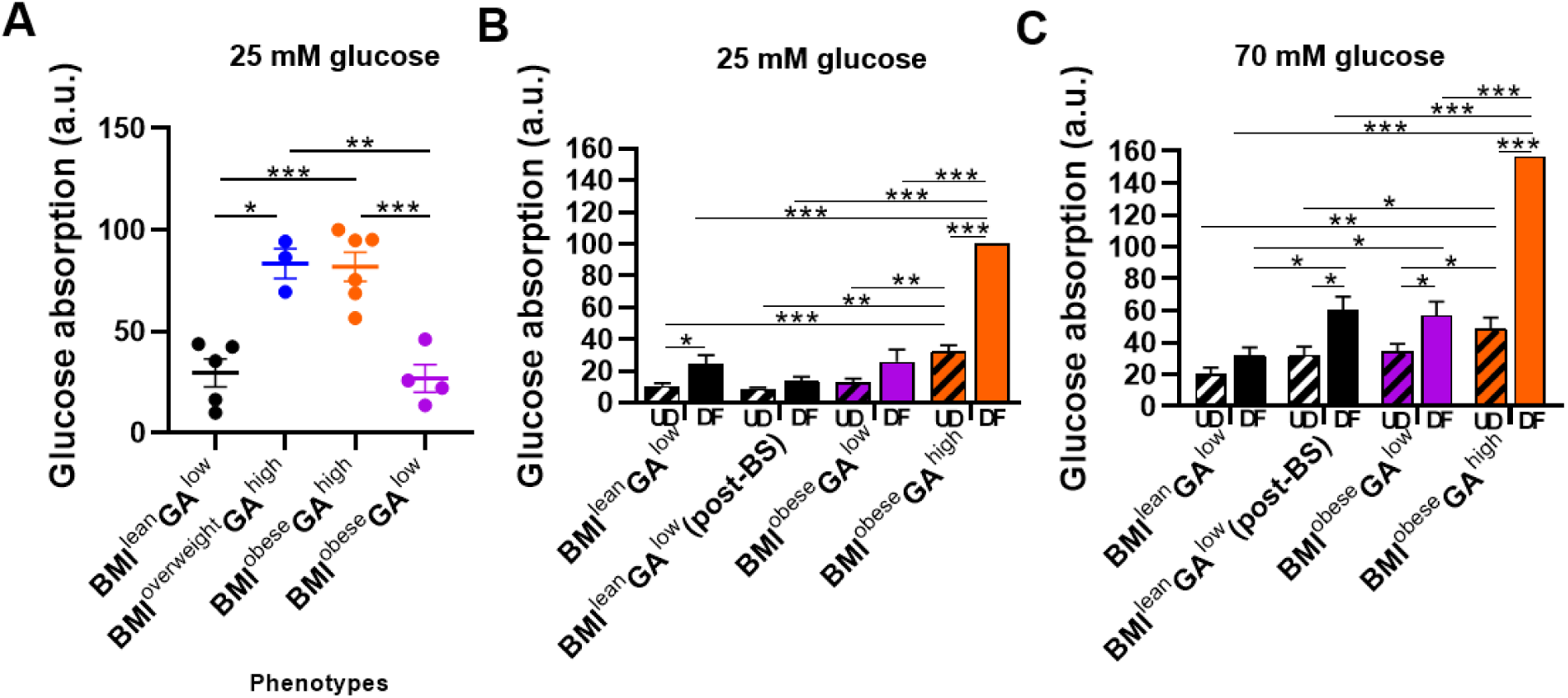
Relationship between subject’s BMI and transepithelial (apical-to-basolateral) glucose absorption in differentiated enteroid monolayers. (A) Glucose absorption in DF-EM treated apically with 25 mM glucose for 120 minutes (twenty-three EM representing eighteen subjects). (B, C) Glucose absorption in UD-EM and DF-EM from four representative enteroid cultures treated apically with 25 mM glucose (B) or 70 mM glucose (C) for 120 minutes. Data in each experiment was normalized to the ‘normalization EM’. Glucose absorption values are provided in Table S2. Data from four-eight independent experiments are presented as mean ± SEM; * p ≤0.05, ** p ≤0.01, *** p ≤0.001 by two-tailed Student’s t-test; ns-not significant (p>0.05) comparisons are not shown. The BMI classification is based on the World Health Organization guidelines. The schematic was created using biorender.com. GA, glucose absorption; BMI, body mass index; UD, undifferentiated; DF, differentiated; EM, enteroid monolayers.

We next compared the physiological and molecular mechanisms of the sugar absorption and metabolism between BMI^lean^ and BMI^obese^ phenotypes using representative enteroid cultures. In order to determine whether BS reduces glucose absorption to levels in naturally lean conditions, enteroids from the BMI^lean^GA^low^ phenotype included a representative culture from each of the two groups: 1) naturally lean or 2) sustainably lean post-BS.

### Changes in luminal glucose concentrations differentially affect glucose absorption in enteroids with BMI^lean^GA^low^ phenotype compared to the BMI^obese^GA^high^ phenotype

We measured glucose absorption in representative DF and UD jejunal enteroids exposed apically to 25 mM glucose (to mimic a low carbohydrate meal). Glucose absorption was higher in DF-EM compared to phenotype matched UD-EM in all groups (Fig.1B, Table S2). The glucose absorption of the BMI^obese^GA^low^ phenotype was similar to the BMI^lean^GA^low^ phenotype. The absorption in DF-EM from BMI^obese^GA^high^ group was significantly higher compared to DF-EM with low glucose absorption. Strikingly, UD-EM from the BMI^obese^GA^high^ phenotype absorbed a significantly higher amount of glucose (∼3 fold) compared to UD-EM from both lean groups, indicating that crypt-based epithelial cells might also contribute to the elevated sugar absorption in obesity. Kinetic analysis shows that the differences in the basolateral glucose concentration between BMI^lean^GA^low^ and BMI^obese^GA^high^ groups were already significant at 30 min after exposure to apical glucose in both DF-EM and UD-EM (Fig. S1A, B; Table S3). We conclude that following treatment with 25 mM glucose, the intestinal epithelium derived from a sub-population of morbidly obese subjects absorbs significantly more glucose than the epithelium derived from lean subjects. Additionally, BMI^obese^GA^low^ phenotype has glucose absorption similar to the BMI^lean^ group, suggesting that obesity is a complex metabolic disease and pathways other than intestinal glucose absorption might be involved.

Next, to mimic the effects of a high carbohydrate meal, 70 mM glucose was added apically to EM. The absorption in DF-EM from BMI^obese^GA^high^ group was significantly higher compared to DF-EM from all three groups with low glucose absorption. The difference was highest (∼5.0 fold) in comparison to the naturally lean BMI^lean^GA^low^ group (Fig. 1B, Table S2). The time course further confirmed significant differences in basolateral glucose concentration between naturally lean BMI^lean^GA^low^ and BMI^obese^GA^high^ groups at all studied time points (Fig. S1C, D; Table S3). Contrary to 25 mM apical glucose, 70 mM apical glucose treatment of DF-EM representing the BMI^obese^GA^low^ phenotype showed significantly higher glucose absorption (∼1.8 fold) compared to the naturally lean BMI^lean^ GA^low^ enteroids (Fig. 1C). EM from BMI^lean^GA^low^ (post-BS) group exposed to high luminal glucose behaved similar to the BMI^obese^GA^low^ group. These data indicate that post-BS epithelium in lean subjects differs from epithelium from naturally lean subjects in which intestinal epithelial glucose absorption remains low regardless of luminal glucose concentrations.

We conclude that the glucose absorption in enteroids from BMI^obese^GA^high^ phenotype correlates with the increase of luminal dietary glucose concentration and appears to not reach a saturation. Importantly, at high apical glucose concentration the glucose absorption by UD-EM from BMI^obese^ GA^high^ group was similar to glucose absorption by DF-EM from BMI^lean^GA^low^ group, further suggesting the contribution of crypt-like epithelium to high serosal glucose in obesity.

In order to test whether elevated intestinal glucose absorption in obesity is preserved in enteroid cultures, the measurements in Fig. 1 were conducted at low passages (11-15) and high passages (30-35) revealing similar trends of glucose absorption in enteroids from all tested phenotypes over the indicated passages. The high rate of concordance in the transcriptome of intestinal organoids and intestinal primary tissue has been recently shown [37, 38], indicating that intestinal organoid cultures preserve the donor phenotype. Based on these published observations, our data suggest that measured differences in epithelial glucose handling between enteroid cultures representing lean or obese phenotypes reflect the intrinsic trait of the donor intestinal epithelium. Overall, our results indicate that enteroids are a faithful and reliable patient-specific model to study obesity-imposed intestinal epithelial pathologies.

### Expression levels of transporters involved in dietary carbohydrate absorption are significantly altered in enteroids from BMI^obese^GA^high^ phenotype compared to BMI^lean^GA^low^ phenotype

To understand the mechanisms responsible for the increased glucose absorption in obesity, we focused on the expression and localization of major carbohydrate transporters SGLT1, GLUT2, GLUT5 and GLUT1 in representative enteroids (one enteroid culture/phenotype) from BMI^obese^GA^high^ and BMI^lean^GA^low^ phenotypes using immunoblotting, immunofluorescence or qRT-PCR (in the absence of reliable antibody).

SGLT1 expression was significantly upregulated in all groups upon differentiation, which is consistent with physiological data that villus enterocytes are mainly involved in dietary glucose absorption. Interestingly, the magnitude of SGLT1 upregulation due to differentiation was significantly higher (∼4 fold) in the BMI^obese^GA^high^ group compared to from the BMI^lean^GA^low^ phenotype (∼2.5 fold) (Fig. 2A, B). Immunofluorescence (Fig. 2C) confirmed that SGLT1, a BBM protein [39], co-localizes with F-actin at the apical membrane in our human EM (Fig. S2). The fluorescence intensity of SGLT1 was higher in EM from BMI^obese^GA^high^ compared to BMI^lean^GA^low^ (Fig. 2C). These data support our glucose transport findings demonstrating a link between increased expression of SGLT1 and increased glucose absorption in BMI^obese^GA^high^ enteroids.

**Fig. 2.**
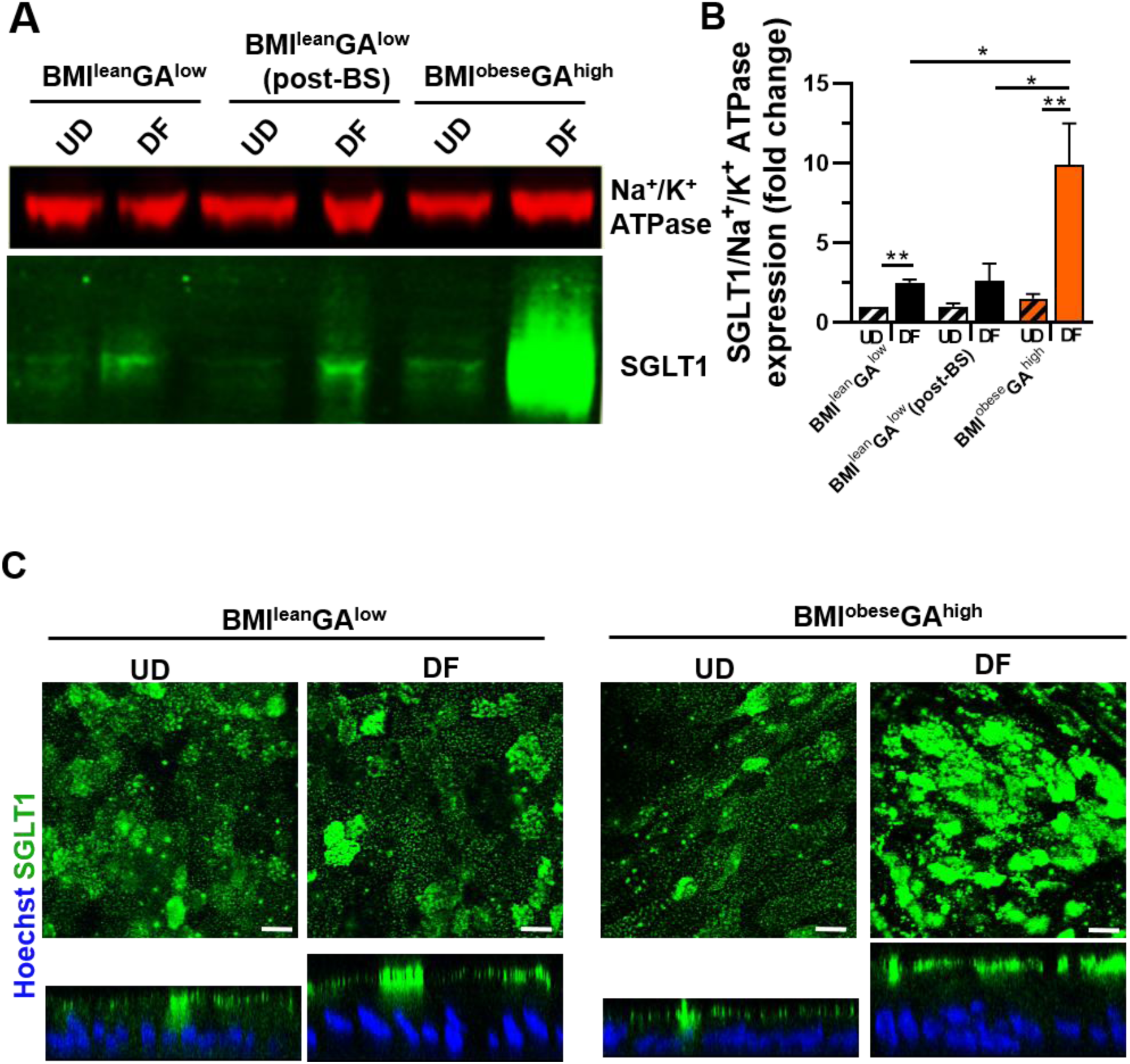
SGLT1 is significantly upregulated in enteroid cultures derived from BMI^obese^GA^high^ phenotype compared to BMI^lean^GA^low^ phenotype. (A) Representative immunoblot of SGLT1 (∼ 77 kDa) and membrane marker Na^+^/K^+^ ATPase-α subunit (∼110 kDa) used as a loading control in total cell membrane lysates. (B) Densitometry quantification of SGLT1 protein expression. Data from three independent experiments (mean ± SEM); * p ≤0.05, ** p ≤0.01 (two-tailed Student’s t-test); ns-not significant (p>0.05) comparisons are not shown. (C) Representative immunofluorescence confocal images (XY and XZ) of EM immunostained for SGLT1 (green). Blue-Hoechst-stained nuclei. Images are representative of three independent experiments. Scale bar, 10 μm. Note that differentiated cells are taller compared to undifferentiated cells in enteroid monolayers. GA, glucose absorption; BMI, body mass index; UD, undifferentiated; DF, differentiated; EM, enteroid monolayers.

Similar to a recent report [7], we also observed that GLUT1 in expressed in human enterocytes (Fig. 3). The localization and relative amount of GLUT1 protein decreased with differentiation. Immunofluorescence demonstrated that GLUT1 was mainly localized to the lateral membrane and intracellularly in UD-EM, and intracellularly in DF-EM from the BMI^lean^GA^low^ group (Fig. 3C, Fig. S3). Interestingly, both GLUT1 expression levels and intracellular distribution were greatly diminished in BMI^obese^GA^high^ enteroids (Fig. 3).

**Fig. 3.**
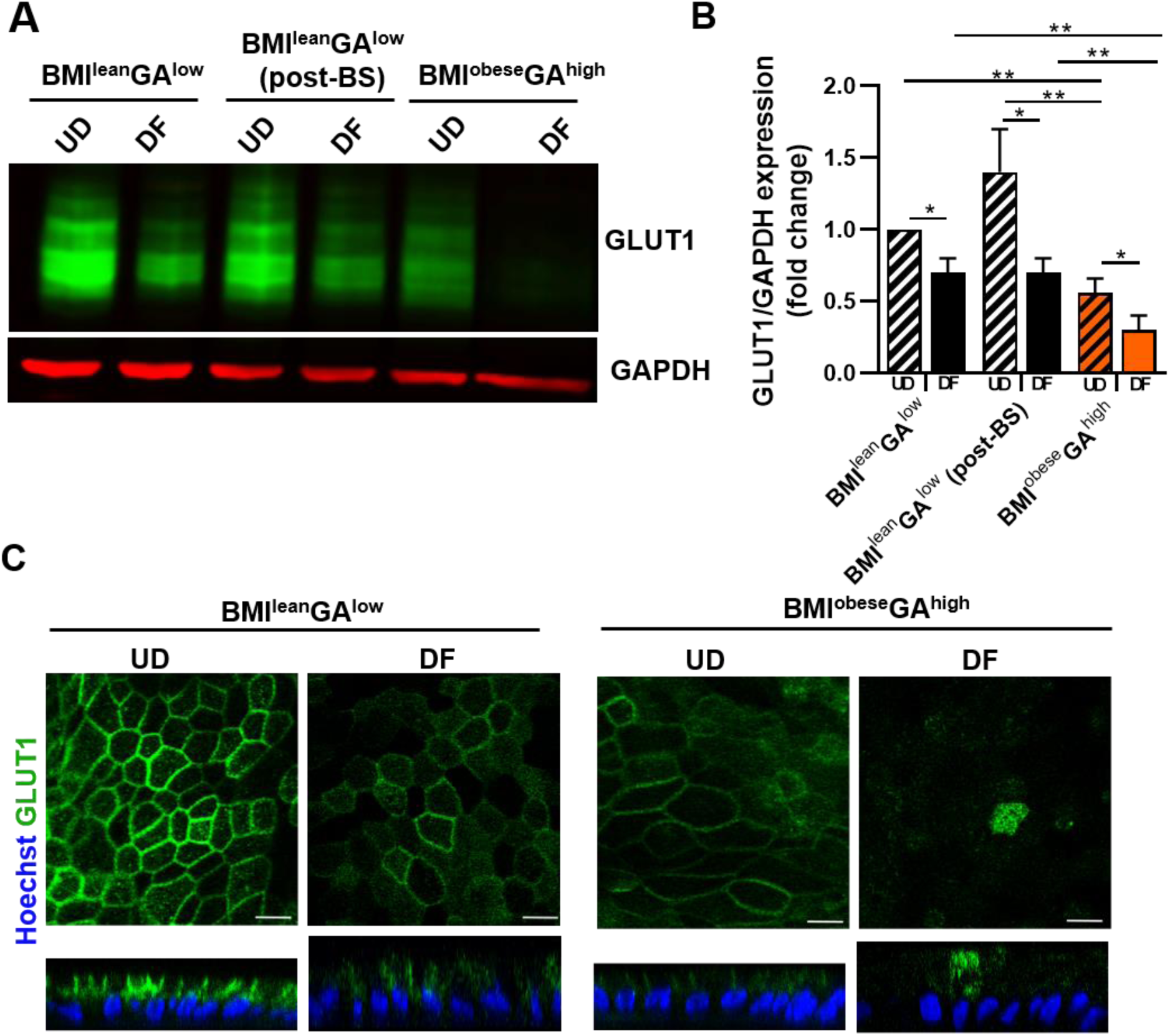
GLUT1 is significantly downregulated in enteroid cultures derived from BMI^obese^GA^high^ phenotype compared to BMI^lean^GA^low^ phenotype. (A) Representative immunoblot of GLUT1 (∼ 55 kDa) and housekeeping marker GAPDH (∼36 kDa) in total cell lysates. (B) Densitometry quantification of GLUT1 protein expression. Data from four independent experiments (mean ± SEM); * p ≤0.05, ** p ≤0.01 (two-tailed Student’s t-test); ns-not significant (p>0.05) comparisons are not shown. (C) Representative immunofluorescence confocal images (XY and XZ) of EM immunostained for GLUT1 (green). Blue-Hoechst-stained nuclei. Images are representative of three independent experiments. Scale bar, 10 μm. Note that differentiated cells are taller compared to undifferentiated cells in enteroid monolayers. GA, glucose absorption; BMI, body mass index; UD, undifferentiated; DF, differentiated; EM, enteroid monolayers.

Similar to SGLT1, *GLUT2* mRNA expression was similar in UD-EM and was significantly increased in DF-EM in all three groups, since dietary glucose absorption is mostly accomplished by villus enterocytes (Fig. 4A). DF-EM representing the BMI^obese^GA^high^ phenotype had higher *GLUT2* mRNA expression (∼55 fold) compared to those from naturally lean BMI^lean^GA^low^ group. This data together with the functional assessment of the contribution of GLUT2 to glucose absorption using phloretin indicates that GLUT2 significantly contributes to the elevated glucose absorption in obesity.

**Fig. 4.**
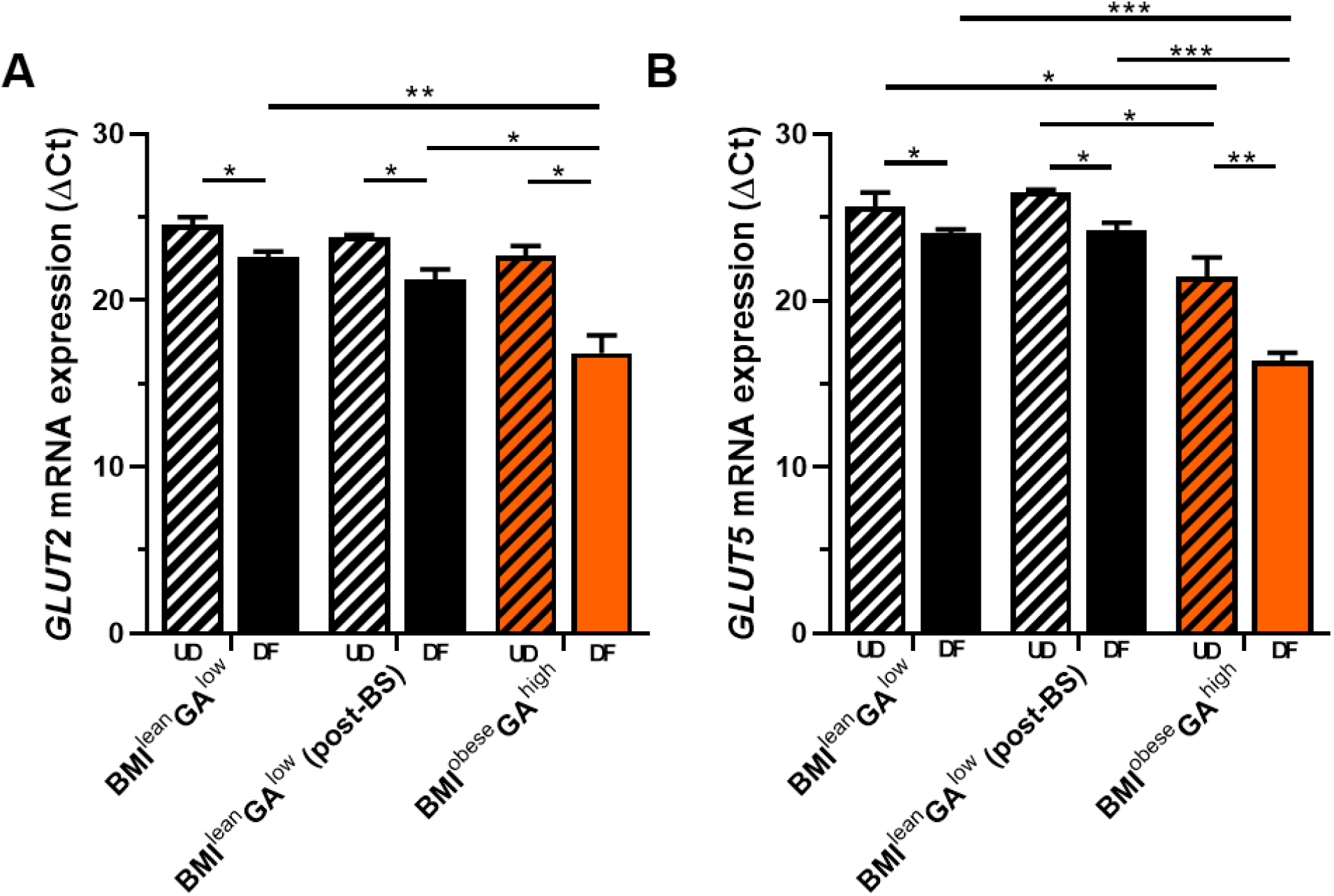
*GLUT2* and *GLUT5* mRNA expression levels are significantly upregulated in enteroid cultures derived from BMI^obese^GA^high^ phenotype compared to BMI^lean^GA^low^ phenotype. qPCR analysis of *GLUT2* mRNA (A) and *GLUT5* mRNA (B) expression in UD and DF enteroid cultures. Data is represented as ΔCt normalized to RNA18S mRNA expression. Data from three independent experiments (mean ± SEM); * p ≤0.05, ** p ≤0.01, *** p ≤0.001 (two-tailed Student’s t-test); ns-not significant (p>0.05) comparisons are not shown. ΔCt values are provided in Table S4. GA, glucose absorption; BMI, body mass index; UD, undifferentiated; DF, differentiated; EM, enteroid monolayers; Ct, cycle threshold.

To determine whether fructose absorption is affected, the mRNA levels of the fructose transporter, *GLUT5*, were measured. *GLUT5* mRNA expression was similar in UD-EM in BMI^lean^GA^low^ and BMI^lean^GA^low^ (post-BS) cultures but was significantly higher (∼20 fold) in the UD-EM of the BMI^obese^GA^high^ phenotype. *GLUT5* mRNA expression significantly increased upon differentiation, and BMI^obese^GA^high^ DF-EM had significantly higher GLUT5 mRNA expression (∼9.5 fold) compared to naturally lean enteroids (Fig. 4B). These data, together with the GLUT5 inhibition data, suggest that fructose absorption may also be higher in EM from BMI^obese^GA^high^ phenotype compared to the lean phenotype.

Taken together, our data indicate that in BMI^obese^ GA^high^ enteroids, obesity is associated with significant changes in expression and/or localization of the major carbohydrate transporting proteins SGLT1, GLUT1, GLUT2 and GLUT5. Importantly, human enteroids in long-term cultures preserve these obesity-related differences in the expression of carbohydrate transporters.

### Inhibition of intestinal sugar transporters SGLT1 and GLUT2 significantly decreases glucose absorption in enteroid monolayers from BMI^obese^GA^high^ phenotype

Both the increase in luminal dietary glucose uptake and basolateral secretion by enterocytes may be responsible for the elevated absorption of glucose reaching the bloodstream. We next examined the contribution of glucose transporters SGLT1, GLUT2, and GLUT1 to overall glucose absorption by pharmacological studies. DF-EM (highest glucose absorption as shown in Fig.1) were exposed to either 25 mM (Fig. 5A) or 70 mM glucose (Fig. 5B, C) in the presence or absence of phloretin (PT, GLUT2 inhibitor) or phloridzin (PZ, SGLT1 inhibitor) [16, 40, 41]. Either PT (50 µM) or PZ (50 µM) treatments significantly decreased glucose absorption compared to untreated controls. The combination of PT and PZ using low (50 µM) or high (400 µM) concentrations did not further decrease the basolateral glucose concentration (Fig. 5, Table S5), indicating that both SGLT1 and GLUT2 are involved in the same pathway for glucose absorption. In contrast, STF-31 (GLUT1 inhibitor) [42] did not affect glucose absorption in EM from the BMI^lean^GA^low^ group (highest expression of GLUT1) (Fig. S4).

**Fig. 5.**
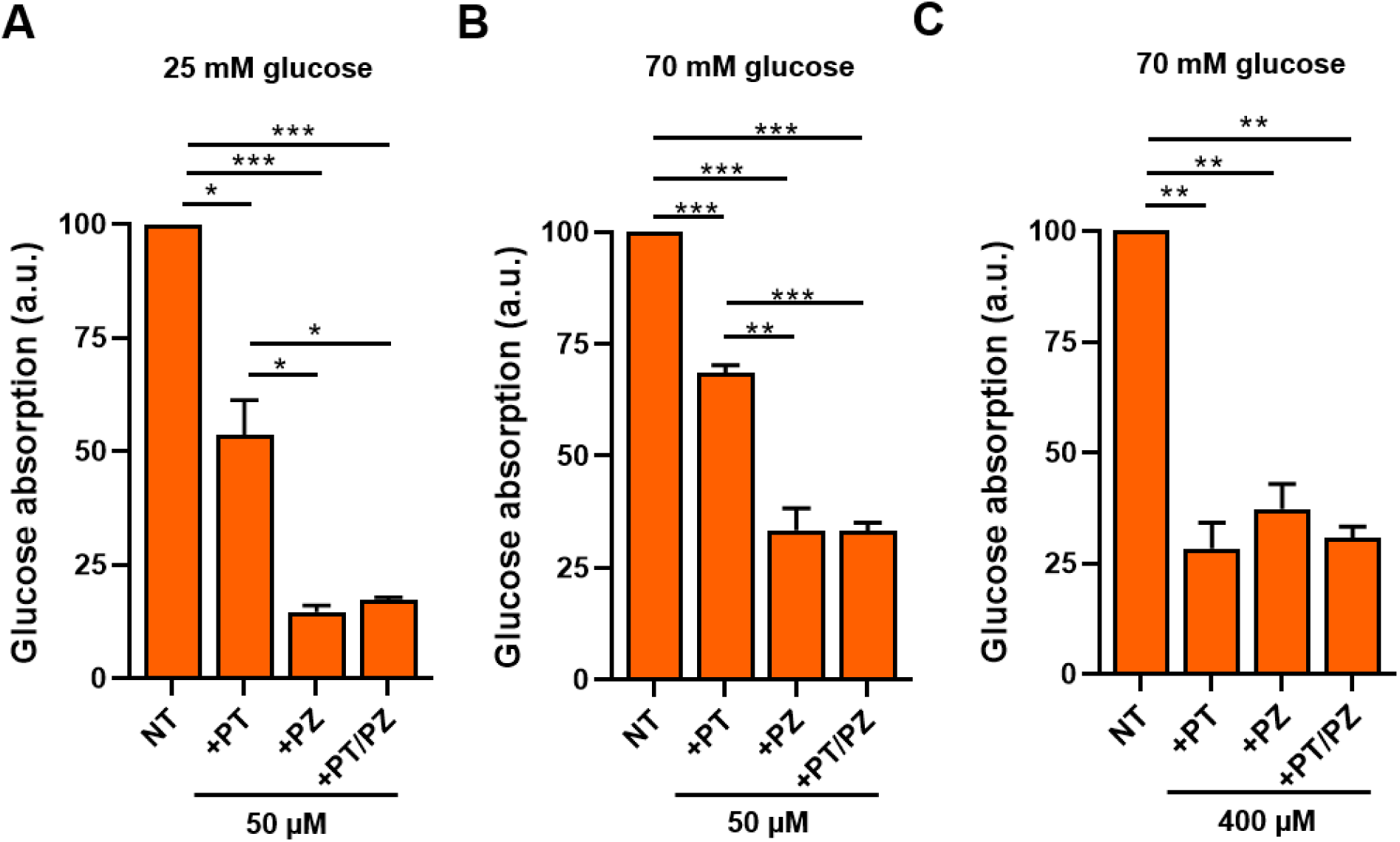
Inhibition of either SGLT1 or GLUT2 significantly decreases glucose absorption in differentiated enteroid cultures derived from BMI^obese^GA^high^ phenotype. Effects of phloridzin (PZ, SGLT1 inhibitor), phloretin (PT, GLUT2 inhibitor), or their combination on glucose absorption in DF-EM derived from BMI^obese^GA^high^ phenotype at 120 min of exposure to either 25 mM (A) or 70 mM (B, C) apical glucose. Basolateral glucose concentration in each experiment was normalized to the value of the NT sample. Data from three independent experiments (mean ± SEM); * p ≤0.05, ** p ≤0.01, *** p ≤0.001 (two-tailed Student’s t-test); ns-not significant (p>0.05) comparisons are not shown. Glucose absorption values are provided in Table S5. NT, not treated; GA, glucose absorption; BMI, body mass index; UD, undifferentiated; DF, differentiated; EM, enteroid monolayers

In order to interrogate whether increased paracellular permeability contributes to the increased glucose absorption observed in the BMI^obese^GA^high^ group, we exposed DF-EM apically with lucifer yellow, the low molecular weight paracellular flux tracer. The fluorescence levels of lucifer yellow in the basolateral media were similar (Fig. S5), indicating that differences in glucose absorption between the BMI^lean^GA^low^ and BMI^obese^GA^high^ groups were based on transcellular transport and not due to glucose leak across tight junctions. Collectively these results suggest that SGLT1 and GLUT2, are the major contributors to increased glucose absorption in obesity. These data are in agreement with the increased expression of SGLT1 and GLUT2 detected in cultures from BMI^obese^GA^high^ phenotype.

### Elevated gluconeogenesis contributes significantly to the increase in serosal glucose in obesity

The expression of the rate-limiting gluconeogenic enzymes, phosphoenolpyruvate carboxykinase 1 (PEPCK1) and glucose-6-phosphatase (G6Pase), in human enterocytes indicates that the proximal small intestine might contribute to *de novo* glucose synthesis [43]. We found that PEPCK1 protein expression was significantly upregulated in DF-EM (∼55 fold) and UD-EM (∼20 fold) from the BMI^obese^GA^high^ group compared to lean enteroids (Fig. 6A, B). Also, *G6Pase* mRNA levels were significantly increased (∼16-fold) in DF-EM from the BMI^obese^GA^high^ phenotype compared to lean cultures (Fig. 6C). The increased expression levels of gluconeogenic enzymes indicate that gluconeogenesis could also contribute to high blood glucose in obesity.

**Fig. 6.**
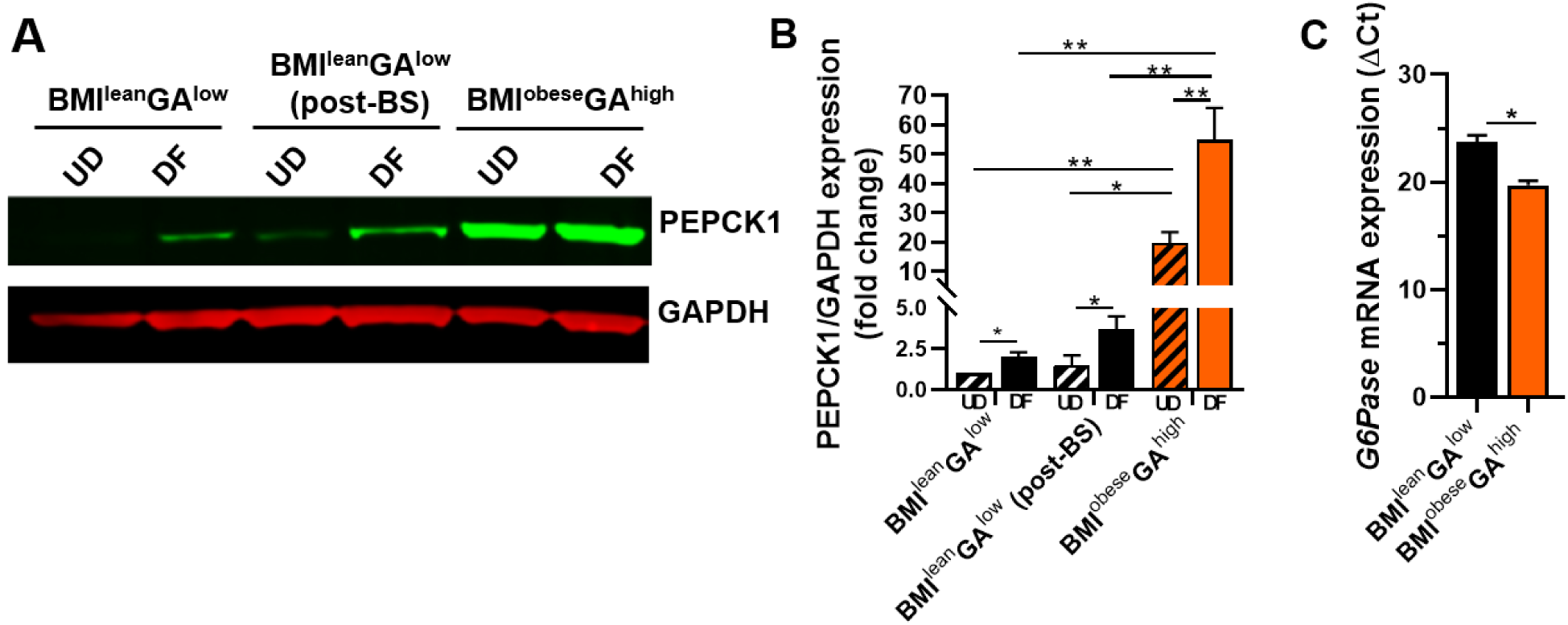
Gluconeogenic enzymes are significantly upregulated in enteroid cultures derived from BMI^obese^GA^high^ phenotype compared to BMI^lean^GA^low^ phenotype. (A) Representative immunoblot of PEPCK1 (∼ 69 kDa) and housekeeping marker GAPDH (∼36 kDa) in total cell lysates. (B) Densitometry quantification of PEPCK1 protein expression. (C) qPCR analysis of glucose-6-phosphatase (G6Pase) mRNA levels in differentiated enteroid cultures derived from BMI^obese^GA^high^ phenotype and BMI^lean^GA^low^ phenotype subjects. Data is represented as ΔCt normalized to RNA18S mRNA expression. Data from three independent experiments (mean ± SEM); * p ≤0.05, ** p ≤0.01 (two-tailed Student’s t-test); ns-not significant (p>0.05) comparisons are not shown. GA, glucose absorption; BMI, body mass index; UD, undifferentiated; DF, differentiated; EM, enteroid monolayers; Ct, cycle threshold.

To test whether the elevated gluconeogenic enzymes increase glucose production by enterocytes in obesity, we incubated DF-EM from representative cultures of BMI^lean^GA^low^ and BMI^obese^GA^high^ groups in glucose-free solutions containing gluconeogenic substrates and measured the amount of glucose deposited into the basolateral media. Apical exposure of DF-EM to the mixture of 2 mM pyruvate/20 mM lactate significantly increased the basolateral glucose concentration in BMI^obese^GA^high^ EM, but not in lean enteroids, in which the basolateral glucose concentration was below detection levels (Fig. 7A, Table S5). These data indicate that *de novo* glucose synthesis via elevated expression of gluconeogenic enzymes may also contribute to increased serosal glucose levels in obesity.

**Fig. 7.**
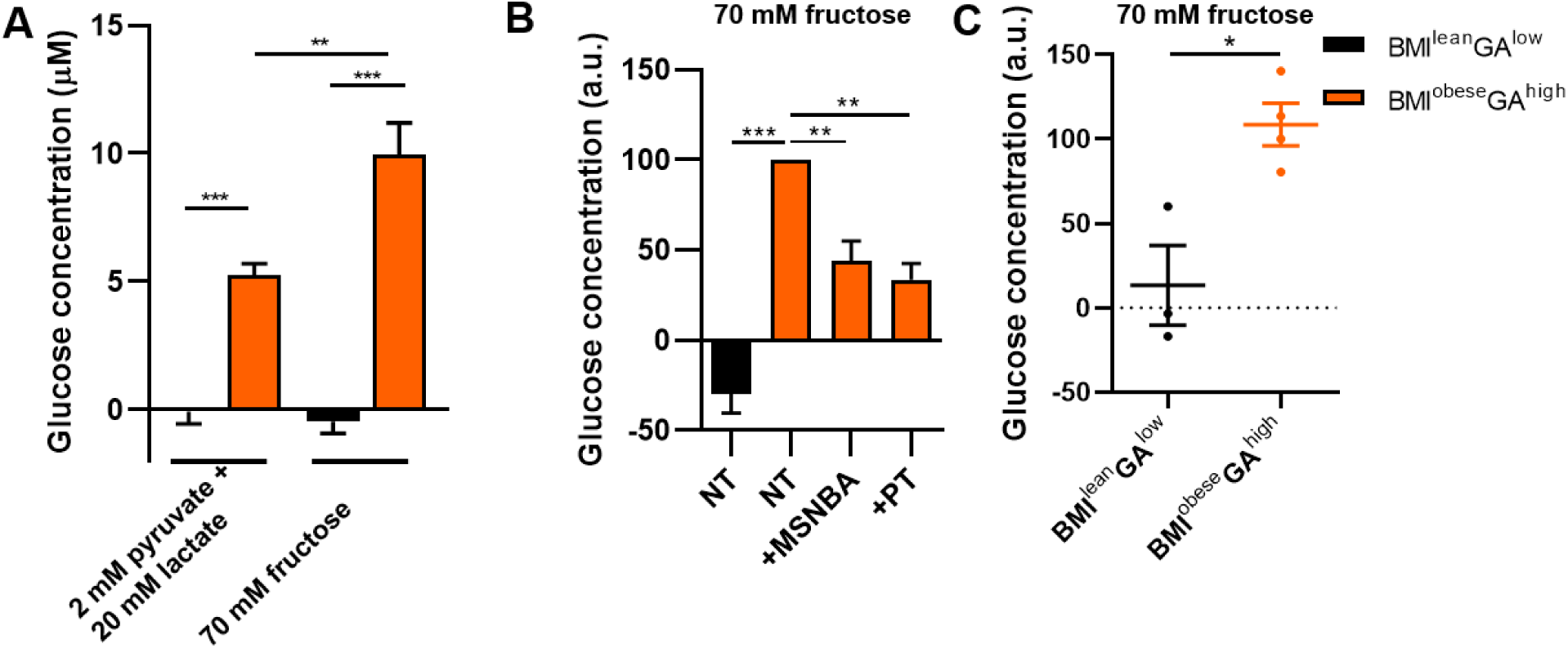
Enteroid monolayers representing the BMI^obese^GA^high^ demonstrate significantly elevated gluconeogenesis with dietary fructose as a potential substrate. (A) Basolateral glucose concentration after apical treatment of DF-EM from BMI^obese^GA^high^ phenotype or BMI^lean^GA^low^ phenotype with either 2 mM pyruvate/20 mM lactate or 70 mM fructose for 3 h. Basolateral glucose concentration in EM representing the BMI^lean^GA^low^ phenotype was below detection limit. Glucose absorption values are provided in Table S5. (B) Effects of 500 µM N-[4- (methylsulfonyl)-2-nitrophenyl]-1,3-benzodioxol-5-amine (MSNBA, GLUT5 inhibitor) or 400 µM phloretin (PT, GLUT2 inhibitor) on basolateral glucose concentration in DF-EM derived from BMI^obese^GA^high^ phenotype at 3 h of exposure to 70 mM apical fructose. Basolateral glucose concentration in each experiment was normalized to the value of the NT sample. Basolateral glucose concentration in EM representing the BMI^lean^GA^low^ phenotype was below detection limit. Glucose absorption values are provided in Table S6. (C) Basolateral glucose concentration after apical treatment of DF-EM from BMI^obese^GA^high^ phenotype (n=3 subjects) or BMI^lean^GA^low^ phenotype (n=4 subjects) with 70 mM fructose for 16 h. Data in each experiment was normalized to the ‘normalization EM’. (A-C) Data from three independent experiments (mean ± SEM); * p ≤0.05, ** p ≤0.01, *** p ≤0.001 (two-tailed Student’s t-test); ns-not significant (p>0.05) comparisons are not shown. NT, not treated; GA, glucose absorption; BMI, body mass index; UD, undifferentiated; DF, differentiated; EM, enteroid monolayers.

Fructose feeding induces significant increases in expression of intestinal fructolytic and gluconeogenic enzymes in mice [44] and ingested fructose is metabolized in the small intestine via gluconeogenesis [29, 44]. We hypothesize that in BMI^obese^GA^high^ enteroids, fructose uptake via elevated apical GLUT5 followed by increased fructose metabolism via elevated gluconeogenic enzymes might significantly contribute to the elevated serosal glucose. We used apical fructose to assess the possible contribution of this dietary sugar to the basolateral glucose deposit via gluconeogenesis. Exposure to 70 mM apical fructose for 3 h significantly increased the basolateral glucose concentration in DF-EM from BMI^obese^GA^high^, but not in the lean phenotype (Fig.7A, Table S5). These results indicate that in the BMI^obese^GA^high^ group, increased GLUT5 expression leads to increased fructose uptake, which serves as a substrate for *de novo* glucose production.

In order to examine the contribution of GLUT5 and GLUT2 to fructose uptake, we performed pharmacological inhibition studies using N-[4-(methylsulfonyl)-2-nitrophenyl]-1,3-benzodioxol-5-amine (MSNBA, GLUT5 inhibitor) [45] and phloretin (PT, GLUT2 inhibitor). DF-EM from BMI^obese^GA^high^ phenotype were treated apically for 3 h with 70 mM fructose in the presence of 500 µM MSNBA or 400 µM PT. Inhibition of either GLUT2 or GLUT5 significant decreased the basolateral glucose concentration (Fig. 7B), indicating that both GLUT2 and GLUT5 are involved in luminal fructose uptake, and their inhibition decreases basolateral glucose deposition.

We next measured the contribution of gluconeogenesis to the basolateral glucose concentration in DF-EM from BMI^lean^GA^low^ and BMI^obese^GA^high^ phenotypes (Fig. 7C) treated with 70 mM fructose for 16 h. The DF-EM from the BMI^obese^GA^high^ phenotype had significantly higher basolateral glucose concentration compared to the BMI^lean^GA^low^ phenotype. Collectively these data suggest that increased basolateral glucose deposition by gluconeogenesis can also substantially contribute to obesity.

## Discussion

We used human stem cell-derived enteroids as a model of small intestinal epithelium [31, 32, 34, 36] to determine possible pathophysiology of dietary sugar absorption in obesity. Using enteroids derived from eighteen patients with different BMIs, we showed four different glucose absorption phenotypes. We were able to identify two different physiologic traits in morbidly obese patients: low or high epithelial sugar absorption. Enteroids from patients representing the BMI^obese^GA^high^ phenotype support previous findings that have shown increased expression of carbohydrate transporters and increased glucose absorption in obesity [11, 24, 46, 47, 48, 49]. Our findings demonstrate that at equal concentrations of dietary glucose, patients with BMI^obese^GA^high^ phenotype would transport more glucose into the systemic circulation compared to non-obese patients; therefore, intestinal carbohydrate transport plays an important role in obesity and metabolic diseases in this category of patients. These data also provide a possible explanation for the inability of changes in diet to combat morbid obesity [3, 4, 5]. However, our enteroid studies show that this occurs only in subset of obese patients. We found another morbidly obese phenotype with low glucose absorption (BMI^obese^GA^low^), and these findings match the presence of normoglycemic and hyperglycemic morbidly obese subjects [50]. The uptake and metabolism of amino acids and lipids is altered in obesity, suggesting that pathways other than carbohydrate uptake and metabolism may play a significant role in BMI^obese^GA^low^ phenotype patients. Our enteroid data on differential glucose absorption in morbidly obese individuals strongly supports the recent continuous glucose monitoring study that identified several “glucotypes” based on the variability in blood glucose levels [51].

Importantly, all enteroids from the lean group that included cultures from three successful post-BS cases exhibit low glucose absorption similar to that measured in naturally lean subjects. Our data matches the results obtained by comparing human small intestinal samples pre- and post-BS in which SGLT1 protein expression was increased in obese patients compared to lean subjects, while SGLT1 protein levels in post-BS was similar to lean controls [52]. It remains to be determined whether success of BS could be predicted by high intestinal glucose absorption prior to surgery.

The physiologic differences between enteroids from BMI^lean^GA^low^ and BMI^obese^GA^high^ phenotypes are at least partially due to significant differences in the expression of intestinal sugar transporters. SGLT1 and GLUT2 are important in intestinal glucose absorption, and our data show that both SGLT1 and GLUT2 are upregulated in enteroid cultures from the BMI^obese^GA^high^ phenotype. Inhibition of either SGLT1 or GLUT2 significantly decreases glucose absorption, indicating that these transporters might be potential targets for hyperglycemia in the BMI^obese^GA^high^ subset of obese patients. Recent clinical studies using licogliflozin, a dual SGLT1/2 inhibitor that targets both the intestinal and renal glucose absorption, have shown promising results in decreasing blood glucose and body weight of obese patients [53].

GLUT5 is an apical fructose transporter and its overexpressed in the intestine of T2D patients [21]. Our data show for the first time the functional link between elevated expression of GLUT5 in human intestinal epithelium and upregulation of the major rate-limiting gluconeogenic enzymes, PEPCK1 and G6Pase, which resulted in a significant increase in glucose deposited into serosa following dietary fructose uptake. Pharmacologic inhibition of GLUT5 decreased glucose production via gluconeogenesis, suggesting that GLUT5 may also be a target in obesity management.

Collectively, our data demonstrate that intestinal glucose absorption via transport and gluconeogenesis might significantly contribute to high blood glucose levels in a subset of morbidly obese patients (Figure 8). Based on these data, we suggest that the decrease in serum glucose concentrations that occurs after BS [9, 10] might be due to minimization of the intestinal contribution to the overall blood glucose levels. Thus, targeting molecular steps involved in intestinal glucose transport and gluconeogenesis may represent a pathway for development of novel drugs to treat metabolic diseases. These data also suggest that multiple molecular targets must be considered when using therapeutic approaches in order to duplicate the effects of BS on the instant decrease of blood glucose levels.

**Fig. 8.**
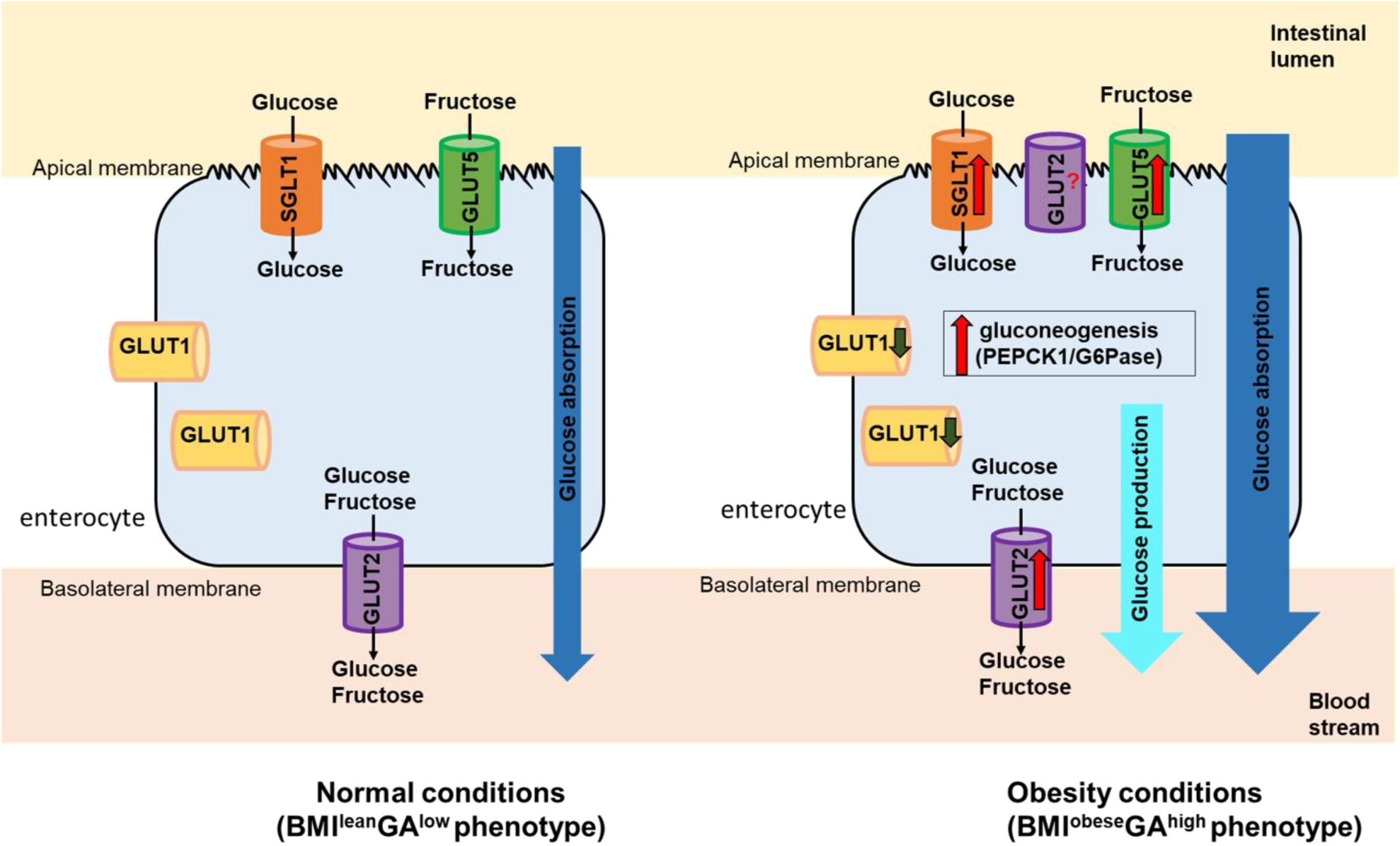
Schematic diagram of glucose transporters, glucose absorption and gluconeogenesis in enterocytes under normal and obesity conditions. Under normal conditions SGLT1 and GLUT5 [17] are localized to the apical membrane of enterocytes transporting glucose and fructose respectively into the cells. GLUT2 is localized at the basolateral membrane and transports glucose and fructose out of the cells into the blood stream [14, 15]. GLUT1 has lateral and intracellular localization. In obesity conditions, expression levels of SGLT1 and GLUT2 are increased resulting in increased glucose absorption. GLUT1 levels are decreased. Additionally, GLUT5 levels and gluconeogenesis enzymes (PEPCK1 and G6Pase) are increased resulting in increased production of glucose in enterocytes that is transported to the blood stream. GLUT2 might be localized apically [24, 25].

Our data confirm observations [7] that the “non-intestinal” GLUT1 glucose transporter is expressed in human enterocytes. Differentiation decreased GLUT1 protein levels and increased its intracellular presence, indicating potential trafficking to the cytosol. Thioredoxin-interacting protein (TXNIP) has a role in GLUT1 endocytosis and degradation of GLUT1 [54]. We observed increase of thioredoxin-interacting protein levels upon differentiation in enteroids (data not shown), suggesting a potential explanation for our observed decreased GLUT1 levels upon differentiation. GLUT1 was not associated with glucose absorption, suggesting a different role in enterocytes. GLUT1 has been shown to increase the supply of glucose to proliferative cancer cells in response to hypoxia; thus, GLUT1 might be supplying glucose to proliferating stem cells in undifferentiated enteroids, which require a lot of energy but display low levels of other sugar transporters and gluconeogenic enzymes. A defined role of GLUT1 in the intestinal epithelium remains to be examined.

Human small intestinal enteroid cultures obtained from different patients can retain the donor phenotype [55]. These data support our findings that human enteroid cultures preserve the phenotypic differences between epithelia from obese, lean and post-BS subjects for at least 42 passages. These findings indicate that stable long-term changes in gene regulation in intestinal stem cells occur in obesity. Enteroids retain these patient characteristics and could serve as a model to study the cellular and molecular changes associated with obesity. Our results are supported by recent observation that other intestinal chronic conditions, as inflammation [37], might also progressively introduce changes to stem cells preserved in organoid cultures and create a disease phenotype that might play an important role in disease progression and a choice of treatment.

We conclude that obesity is characterized by significant and stable changes in the expression and activity of major dietary sugar transporters and gluconeogenic enzymes in human intestinal epithelium in a subset of morbidly obese patients. These changes lead to significant increases in glucose absorption that might substantially contribute to high blood glucose levels in obesity. Human enteroid monolayers, which preserve the patient phenotype in long-term cultures, represent a reliable and robust model to study the consequences of obesity and to search for effective therapeutic interventions.

