## Supplemental Files for "Obesity phenotypes are preserved in intestinal stem cell enteroids from morbidly obese patients"

**Supplementary Information**

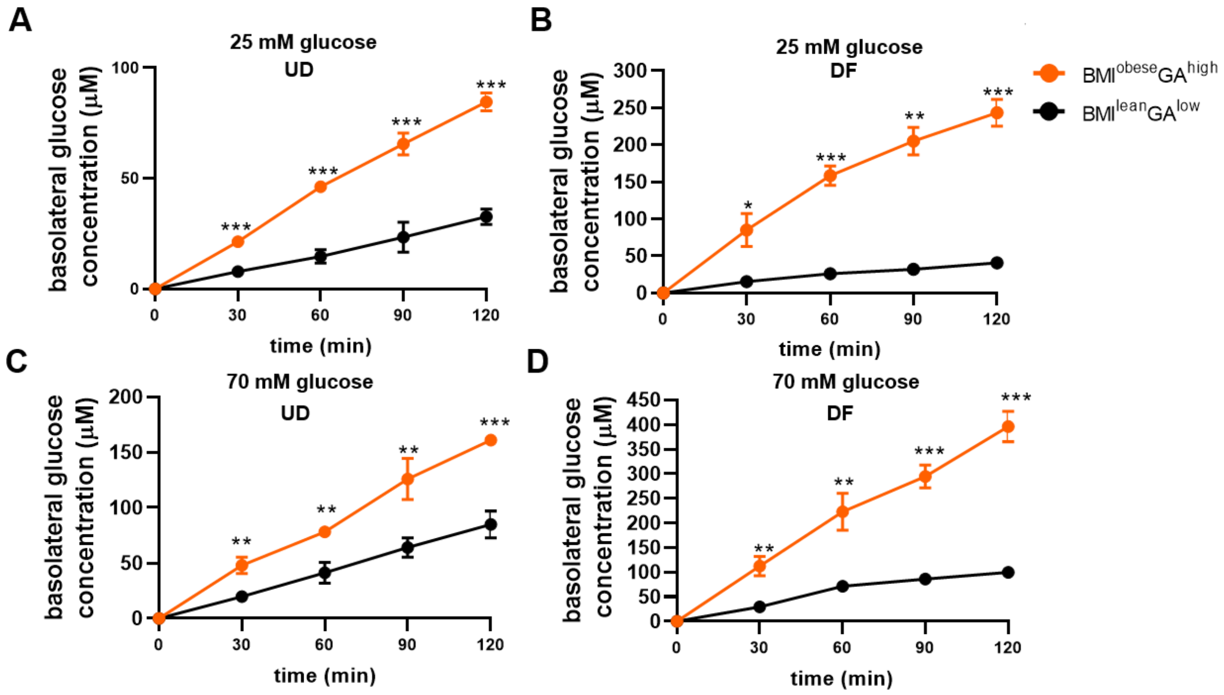

**Fig S1. Enteroid monolayers derived from BMI$^{obese}$GA$^{high}$ phenotype absorb significantly more dietary glucose from lumen to serosa compared to enteroids derived from BMI$^{lean}$GA$^{low}$ phenotype. (**A to D) Time course of transepithelial (apical-to-basolateral) glucose absorption in undifferentiated (A, C) and differentiated (B, D) enteroid monolayers treated apically with either 25 mM (A, B) or 70 mM glucose (C, D). Data from three-six independent experiments are presented as mean ± SEM.; ns-not significant ($p>0.05$), * $p \leq 0.05$, ** $p \leq 0.01$, *** $p \leq 0.001$, **** $p \leq 0.0001$ (two-tailed Student's t-test). GA, glucose absorption; BMI, body mass index; UD, undifferentiated; DF, differentiated; EM, enteroid monolayers.

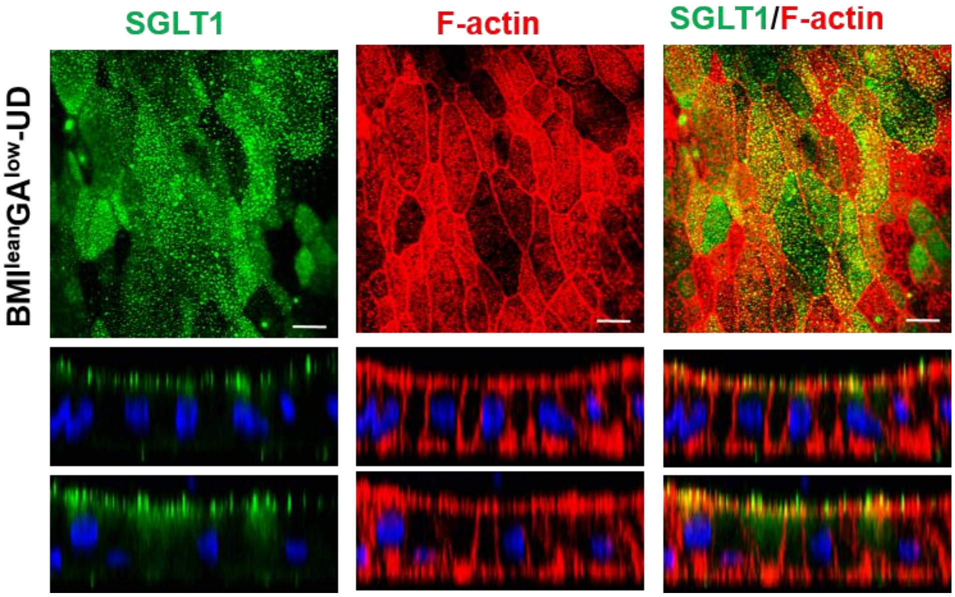

**Fig. S2. SGLT1 is localized at the brush border membrane in enteroid monolayers.** Representative immunofluorescence confocal images (XY and XZ) of UD-DF representing the BMI$^{lean}$GA$^{low}$ phenotype immunostained for SGLT1 (green), F-actin (red) and nuclei (blue). Phalloidin was used to detect F-actin. Images are representative of three independent experiments. Scale bar, 10 μm. GA, glucose absorption; BMI, body mass index; DF, differentiated; EM, enteroid monolayers.

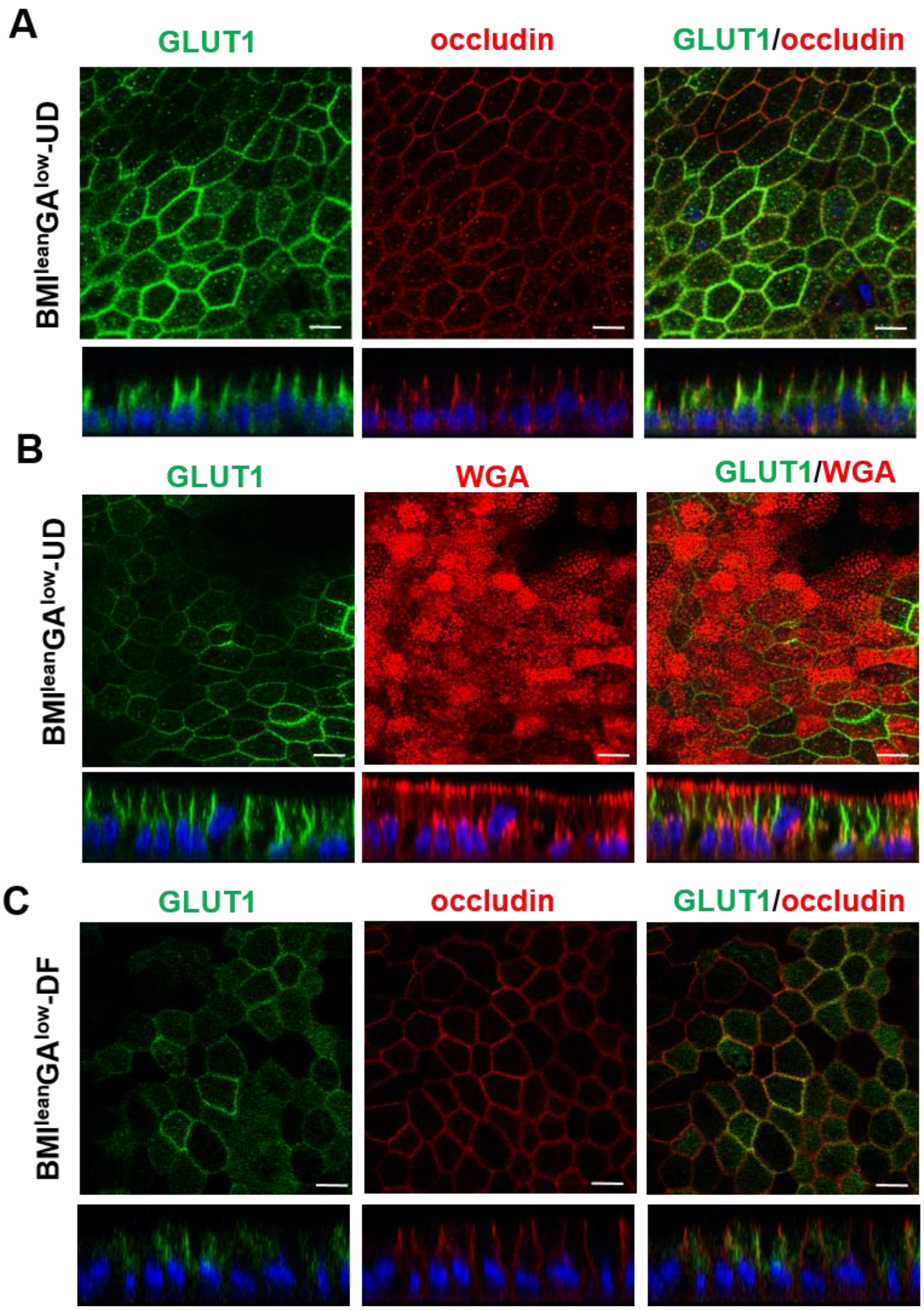

**Fig. S3. GLUT1 is expressed at the lateral membranes in enteroid monolayers and the lateral localization is decreased upon differentiation**. Representative immunofluorescence confocal images (XY and XZ) of UD and DF EM representing the $BMI^{lean}GA^{low}$ phenotype. (A) GLUT1 (green) and occludin (red) immunostaining in UD-EM show lateral expression pattern of GLUT1 just below TJ marked by occludin expression. (B) GLUT1 (green) and WGA (red) immunostaining in UD-EM show that GLUT1 does not appear at the apical surface of enteroids labeled by WGA. **(C)** GLUT1 (green) and occludin (red) immunostaining in DF-EM show lack of colocalization. Blue- nuclear staining by Hoechst. Scale bar, 10 μm. GA, glucose absorption; BMI, body mass index; UD, undifferentiated; DF, differentiated; EM, enteroid monolayers.

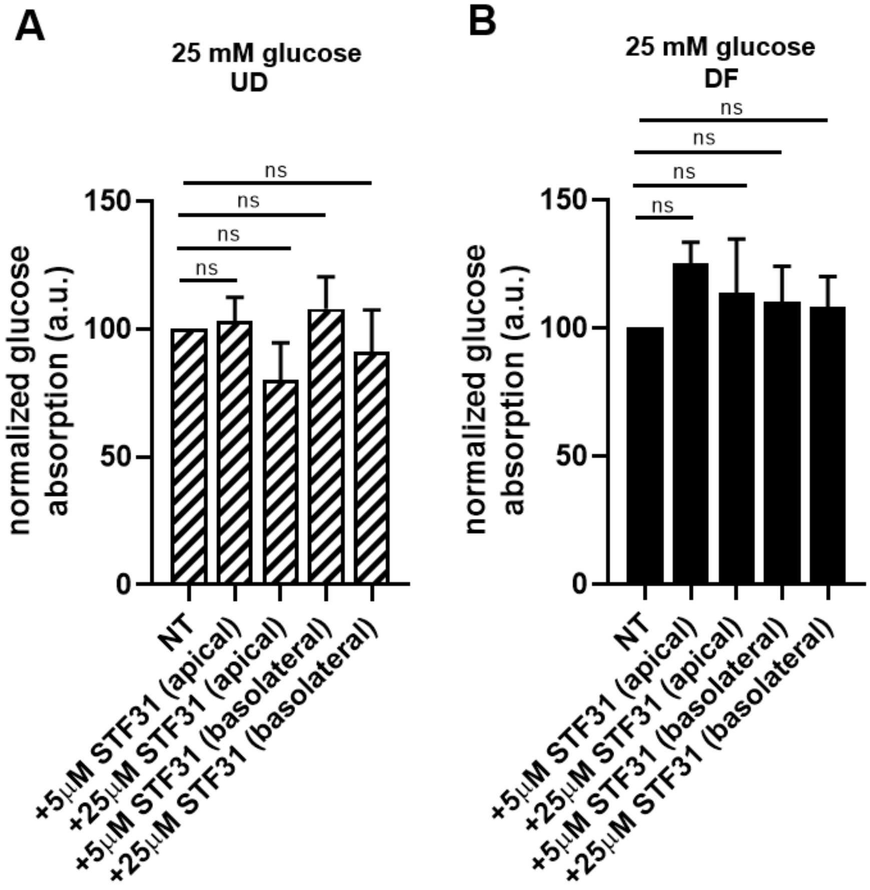

**Fig. S4. GLUT1 inhibition does not have an effect on glucose absorption in enteroids representing the BMI$^{lean}$GA$^{low}$ phenotype.** Normalized basolateral glucose concentration after 120 min of 25 mM apical glucose treatment in the presence of GLUT1 inhibitor STF-31 added either apically or basolaterally (5 µM or 25 µM) in UD (A) DF (B) EM. Basolateral glucose concentration in each experiment was normalized to the value of the NT sample. Data presented as mean ± SEM.; ns-not significant ($p > 0.05$) in comparison with control (two-tailed Student's t-test). NT, not treated; GA, glucose absorption; BMI, body mass index; UD, undifferentiated; DF, differentiated; EM, enteroid monolayers.

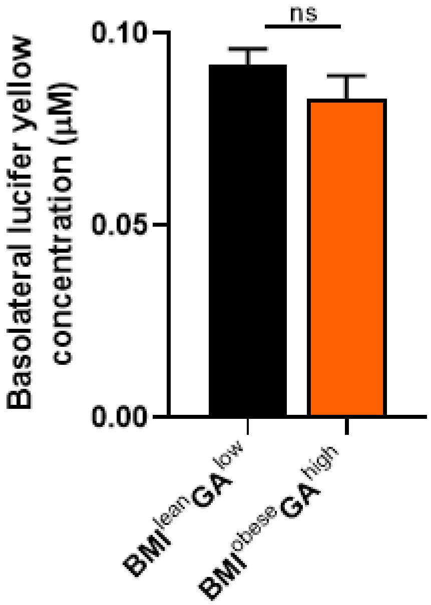

**Fig. S5. Paracellular permeability in enteroid monolayers representing the BMI$^{lean}$GA$^{low}$ and BMI$^{obese}$GA$^{high}$ phenotypes is similar.** DF-EM were treated apically with 200 µM lucifer yellow for 2 h and the fluorescence levels of lucifer yellow in the basolateral media were measured. Data from three independent experiments are presented as mean ± SEM, ns-not significant ($p > 0.05$) in comparison with control (two-tailed Student's t-test). GA, glucose absorption; BMI, body mass index; UD, undifferentiated; DF, differentiated; EM, enteroid monolayers.

### Supplementary Tables

| Short designation | Phenotype | Part of intestine | BMI category | BS status | Gender | Age |
|---|---|---|---|---|---|---|
| Enteroids-1 | $BMI^{lean}GA^{low}$ | Proximal small intestine | lean | none | M | ns |
| Enteroids-2 | $BMI^{lean}GA^{low}$ | Proximal small intestine | lean | none | F | ns |
| Enteroids-3* | $BMI^{lean}GA^{low}$ | Distal small intestine | lean | none | F | ns |
| Enteroids-4 | $BMI^{lean}GA^{low}$ | Distal small intestine | lean | Post VBG | F | ns |
| Enteroids-5* | $BMI^{lean}GA^{low}$ | Proximal small intestine | lean | Post RYGB | F | 46 |
| Enteroids-6 | $BMI^{lean}GA^{low}$ | Proximal small intestine | lean | Post RYGB | F | 43 |
| Enteroids-7 | $BMI^{lean}GA^{low}$ | Distal small intestine | lean | Post RYGB | F | 43 |
| Enteroids-8 | $BMI^{overweight}GA^{high}$ | Proximal small intestine | overweight | Post RYGB | F | 58 |
| Enteroids-9 | $BMI^{overweight}GA^{high}$ | Proximal small intestine | overweight | Post RYGB | F | 51 |
| Enteroids-10 | $BMI^{overweight}GA^{high}$ | Distal small intestine | overweight | Post RYGB | F | 37 |
| Enteroids-11* | $BMI^{obese}GA^{high}$ | Distal small intestine | obese | Pre RYGB | ns | ns |
| Enteroids-12 | $BMI^{obese}GA^{high}$ | Distal small intestine | obese | Pre RYGB | ns | ns |
| Enteroids-13 | $BMI^{obese}GA^{high}$ | Proximal jejunum | obese | Post RYGB | F | 44 |
| Enteroids-14 | $BMI^{obese}GA^{high}$ | Distal jejunum | obese | Post RYGB | F | 44 |
| Enteroids-15 | $BMI^{obese}GA^{high}$ | Proximal small intestine | obese | Post RYGB | F | 56 |
| Enteroids-16 | $BMI^{obese}GA^{high}$ | Proximal small intestine | obese | Post RYGB | F | 67 |
| Enteroids-17 | $BMI^{obese}GA^{high}$ | Distal small intestine | obese | Post RYGB | F | 67 |
| Enteroids-18 | $BMI^{obese}GA^{high}$ | Proximal small intestine | obese | Post RYGB | F | 42 |
| Enteroids-19 | $BMI^{obese}GA^{high}$ | Distal small intestine | obese | Post RYGB | F | 42 |
| Enteroids-20* | $BMI^{obese}GA^{low}$ | Distal small intestine | obese | Pre RYGB | ns | ns |
| Enteroids-21 | $BMI^{obese}GA^{low}$ | Proximal small intestine | obese | Post SG | M | 67 |
| Enteroids-22 | $BMI^{obese}GA^{low}$ | Proximal small intestine | obese | Post RYGB | F | 49 |
| Enteroids-23 | $BMI^{obese}GA^{low}$ | Distal small intestine | obese | Pre RYGB | F | 48 |

**Table S1. Clinical data of patient derived enteroids.**

The BMI classification is based on the World Health Organization guidelines: lean (BMI <25 kg/m$^2$), overweight (BMI 25-30 kg/m$^2$) or obese (BMI> 35 kg/m$^2$).

Abbreviations:

Phenotype; BMI: body mass index GA: glucose absorption
Gender; M: male, F: female
Bariatric surgery (BS) status; VBG: Vertical banded gastroplasty, RYGB: Roux-en-Y Gastric Bypass, SG: sleeve gastrectomy
Gender/Age: ns: not -specified (clinical data not collected)

| | Normalized glucose absorption (mean±SEM) (a.u.) | | | | | | | |
|---|---|---|---|---|---|---|---|---|
| | 25 mM apical glucose | | | | | | | |
| | BMI$^{lean}$GA$^{low}$ | | BMI$^{lean}$GA$^{low}$ (post-BS) | | BMI$^{obese}$GA$^{low}$ | | BMI$^{obese}$GA$^{high}$ | |
| **Time (min)** | **UD** | **DF** | **UD** | **DF** | **UD** | **DF** | **UD** | **DF** |
| **0** | 0.0 ± 0.0 | 0.0 ± 0.0 | 0.0 ± 0.0 | 0.0 ± 0.0 | 0.0 ± 0.0 | 0.0 ± 0.0 | 0.0 ± 0.0 | 0.0 ± 0.0 |
| **30** | 2.3 ± 0.5 | 5.2 ± 1.4 | 1.8 ± 0.1 | 5.3 ± 1.9 | 3.2 ± 0.6 | 7.4 ±4.5 | 8.0 ± 1.1 | 26.3 ± 4.2 |
| **60** | 4.5 ± 0.9 | 11.8 ± 3.2 | 3.6 ± 0.1 | 6.2 ± 1.7 | 5.9 ± 1.1 | 15.2 ± 7.7 | 15.7 ± 2.2 | 50.7 ± 3.9 |
| **90** | 7.0 ± 1.4 | 17.5 ±5.2 | 5.8 ± 0.3 | 13.3 ± 4.6 | 8.6 ± 1.7 | 19.5 ±8.7 | 23.7 ± 3.1 | 76.7 ± 2.9 |
| **120** | 10.5±1.8 | 24.1±5.9 | 8.9±0.9 | 14.6±2.8 | 13.1±2.1 | 25.8±7.6 | 32.4±3.9 | 100.0±0.0 |

| | Normalized glucose absorption (mean±SEM) (a.u.) | | | | | | | |
|---|---|---|---|---|---|---|---|---|
| | 70 mM apical glucose | | | | | | | |
| | BMI$^{lean}$GA$^{low}$ | | BMI$^{lean}$GA$^{low}$ (post-BS) | | BMI$^{obese}$GA$^{low}$ | | BMI$^{obese}$GA$^{high}$ | |
| **Time (min)** | **UD** | **DF** | **UD** | **DF** | **UD** | **DF** | **UD** | **DF** |
| **0** | 0.0 ± 0.0 | 0.0 ± 0.0 | 0.0 ± 0.0 | 0.0 ± 0.0 | 0.0 ± 0.0 | 0.0 ± 0.0 | 0.0 ± 0.0 | 0.0 ± 0.0 |
| **30** | 5.5 ± 1.0 | 9.3 ± 1.6 | 10.4 ± 1.7 | 16.7 ± 2.5 | 10.2 ± 1.5 | 15.5 ± 2.9 | 15.1 ± 2.2 | 50.9 ± 3.2 |
| **60** | 10.3 ± 1.7 | 19.1 ± 4.1 | 18.1 ± 3.5 | 34.4 ± 5.4 | 18.6 ± 3.0 | 30.5 ± 4.9 | 24.7 ± 3.2 | 89.6 ± 2.0 |
| **90** | 15.9 ± 2.9 | 25.0 ± 5.2 | 24.9 ± 3.9 | 49.2 ± 8.2 | 25.6 ± 4.2 | 43.2 ± 5.3 | 37.5 ± 5.6 | 119.1 ± 2.0 |
| **120** | 20.5±4.0 | 31.5±5.4 | 32.2±5.0 | 60.5±8.4 | 34.3±4.9 | 57.3±8.3 | 48.5±7.1 | 156.9±0.0 |

**Table S2. Normalized basolateral glucose concentration in enteroids representing BMI$^{lean}$GA$^{low}$, BMI$^{obese}$GA$^{low}$, and BMI$^{obese}$GA$^{high}$ phenotypes after apical treatment with 25 mM glucose or 70 mM glucose.** Related to Fig.1. Glucose absorption in representative UD-EM and DF-EM treated apically with 25 mM glucose (upper panel) or 70 mM glucose (lower panel) for up to 120 minutes. Data in each experiment was normalized to the 'normalization EM' at 120 min following 25 mM glucose treatment. Data from three-six independent experiments are presented as mean ± SEM. GA, glucose absorption; BMI, body mass index; UD, undifferentiated; DF, differentiated; EM, enteroid monolayers.

| | Basolateral glucose concentration (mean±SD) (µM) | | | |
|---|---|---|---|---|
| | 25 mM apical glucose | | | |
| | BMI$^{lean}$GA$^{low}$ | | BMI$^{obese}$GA$^{high}$ | |
| **Time (min)** | **UD** | **DF** | **UD** | **DF** |
| **0** | 0.0±0.0 | 0.0±0.0 | 0.0±0.0 | 0.0±0.0 |
| **30** | 7.9±0.6 | 15.5±5.8 | 21.4±1.9 | 85.1±22.5 |
| **60** | 14.7±3.0 | 26.0±4.4 | 46.3±2.2 | 158.5±12.9 |
| **90** | 23.4±6.9 | 32.0±3.3 | 65.6±4.9 | 205.1±17.7 |
| **120** | 32.7±3.5 | 40.7±4.0 | 84.6±4.1 | 243.5±17.9 |

| | Basolateral glucose concentration (mean±SD) (µM) | | | |
|---|---|---|---|---|
| | 70 mM apical glucose | | | |
| | BMI$^{lean}$GA$^{low}$ | | BMI$^{obese}$GA$^{high}$ | |
| **Time (min)** | **UD** | **DF** | **UD** | **DF** |
| **0** | 0.0±0.0 | 0.0±0.0 | 0.0±0.0 | 0.0±0.0 |
| **30** | 19.7±0.6 | 29.3±5.6 | 47.9±7.3 | 112.0±19.9 |
| **60** | 41.3±9.3 | 71.1±4.1 | 78.3±4.6 | 222.9±37.3 |
| **90** | 64.2±8.8 | 86.2±7.1 | 125.9±18.7 | 284.3±40.3 |
| **120** | 74.9±12.0 | 99.8±10.1 | 161.0±5.1 | 395.6±30.8 |

**Table S3. Basolateral glucose concentration in enteroids representing BMI$^{lean}$GA$^{low}$ and BMI$^{obese}$GA$^{high}$ phenotypes after apical treatment with 25 mM glucose or 70 mM glucose.** Related to Fig. S1A-D. Glucose absorption in representative UD-EM and DF-EM treated apically with 25 mM glucose (upper panel) or 70 mM glucose (lower panel) for up to 120 minutes. Data from a representative experiment are presented as mean ± SD. GA, glucose absorption; BMI, body mass index; UD, undifferentiated; DF, differentiated; EM, enteroid monolayers.

| | | Relative expression ($\Delta$Ct) (mean ± SEM) |
|---|---|---|
| ***GLUT2*** | | |
| BMI$^{lean}$GA$^{low}$ | UD | 24.5 ± 0.5 |
| BMI$^{lean}$GA$^{low}$ | DF | 22.6 ± 0.3 |
| BMI$^{lean}$GA$^{low}$ (post-BS) | UD | 23.8 ± 0.1 |
| BMI$^{lean}$GA$^{low}$ (post-BS) | DF | 21.3 ± 0.6 |
| BMI$^{obese}$GA$^{high}$ | UD | 22.7 ± 0.6 |
| BMI$^{obese}$GA$^{high}$ | DF | 16.8 ± 1.1 |
| ***GLUT5*** | | |
| BMI$^{lean}$GA$^{low}$ | UD | 25.8 ± 0.7 |
| BMI$^{lean}$GA$^{low}$ | DF | 24.0 ± 0.3 |
| BMI$^{lean}$GA$^{low}$ (post-BS) | UD | 26.5 ± 0.1 |
| BMI$^{lean}$GA$^{low}$ (post-BS) | DF | 24.2 ± 0.5 |
| BMI$^{obese}$GA$^{high}$ | UD | 21.4 ± 1.2 |
| BMI$^{obese}$GA$^{high}$ | DF | 16.4 ± 0.5 |
| ***G6Pase*** | | |
| BMI$^{lean}$GA$^{low}$ | DF | 23.7 ± 0.6 |
| BMI$^{obese}$GA$^{high}$ | DF | 19.7 ± 0.4 |

**Table S4: mRNA expression data for *GLUT2, GLUT5* and *G6Pase*.** Related to Fig.4 and Fig. 6C. qPCR analysis of *GLUT2*, *GLUT5* and *G6Pase* mRNA expression in enteroid cultures representing the BMI$^{lean}$GA$^{low}$ and BMI$^{obese}$GA$^{high}$ phenotypes. Data from three independent experiments is represented as $\Delta$Ct and normalized to RNA18S mRNA expression (mean ± SEM). GA, glucose absorption; BMI, body mass index; UD, undifferentiated; DF, differentiated; EM, enteroid monolayers; Ct, cycle threshold.

| | **Normalized basolateral glucose concentration (a.u.) (mean ± SEM)** |
|---|---|
| **Apical treatment with 25 mM glucose (2 h)** | |
| Not treated (NT) | 100.0 ± 0.0 |
| 50 µM phloretin (PT) | 536.6 ± 7.7 |
| 50 µM phloridzin (PZ) | 14.6 ± 1.4 |
| 50 µM phloretin/ phloridzin (PT/PZ) | 17.2 ± 0.6 |
| **Apical treatment with 70 mM glucose (2 h)** | |
| Not treated (NT) | 100 ± 0.0 |
| 50 µM phloretin (PT) | 68.5 ± 1.8 |
| 50 µM phloridzin (PZ) | 33.4 ± 4.9 |
| 50 µM phloretin/ phloridzin (PT/PZ) | 33.3 ± 1.8 |
| **Apical treatment with 70 mM glucose (2 h)** | |
| Not treated (NT) | 100.0 ± 0.0 |
| 400 µM phloretin (PT) | 28.4 ± 5.8 |
| 400 µM phloridzin (PZ) | 37.3 ± 5.6 |
| 400 µM phloretin/ phloridzin (PT/PZ) | 30.8 ± 2.5 |

**Table S5. Normalized basolateral glucose concentration in enteroids representing the BMI$^{obese}$GA$^{high}$ phenotype after apical treatment with 25mM glucose or 70 mM glucose in the presence of inhibitors.** Related to Fig.5. Basolateral glucose concentration in differentiated enteroid monolayers derived from obese patients exposed apically for 120 min to either 25 mM or 70 mM glucose in the presence of 50 µM or 400 µM phloridzin (PZ, SGLT1 inhibitor), phloretin (PT, GLUT2 inhibitor), or their combination. Basolateral glucose concentration in each experiment was normalized to the value of the NT sample. Data from three independent experiments are presented as mean ± SEM.

| | **Basolateral glucose concentration (µM) (mean ± SEM)** |
|---|---|
| **Apical treatment with 2 mM pyruvate/ 20 mM lactate (3 h)** | |
| BMI$^{lean}$GA$^{low}$-DF | -0.1 ± 0.5 |
| BMI$^{obese}$GA$^{high}$-DF | 5.2 ± 0.5 |
| **Apical treatment with 70 mM fructose (3 h)** | |
| BMI$^{lean}$GA$^{low}$-DF | -0.5 ± 0.5 |
| BMI$^{obese}$GA$^{high}$-DF | 9.9 ±1.3 |

**Table S6. Basolateral glucose concentration in enteroids representing BMI$^{lean}$GA$^{low}$ and BMI$^{obese}$GA$^{high}$ phenotypes after apical treatment with gluconeogenic substrates.** Related to Fig.7A. Basolateral glucose concentration in differentiated enteroid monolayers exposed apically for 3 h to either 2 mM pyruvate/20 mM lactate or 70 mM fructose. Basolateral glucose concentration in EM representing the BMI$^{lean}$GA$^{low}$ phenotype was below detection limit. Data from three independent experiments are presented as mean ± SEM. GA, glucose absorption; BMI, body mass index; DF, differentiated; EM, enteroid monolayers.

| | **Normalized basolateral glucose concentration (a.u.) (mean ± SEM)** |
|---|---|
| **Apical treatment with 70 mM fructose (3 h)** | |
| **BMI$^{lean}$GA$^{low}$-DF** | |
| Not treated (NT) | -29.6±10.9 |
| **BMI$^{obese}$GA$^{high}$-DF** | |
| Not treated (NT) | 100.0 ± 0.0 |
| 500 µM MSNBA | 43.7 ± 11.2 |
| 400 µM phloretin (PT) | 33.4 ± 9.1 |

**Table S7. Normalized basolateral glucose concentration in enteroids representing BMI$^{lean}$GA$^{low}$ and BMI$^{obese}$GA$^{high}$ phenotypes after apical treatment with 70 mM fructose in the presence of inhibitors.** Related to Fig.7B. Basolateral glucose concentration in differentiated enteroid monolayers exposed apically for 3h min to 70 mM fructose in the presence of 500 µM N-[4-(methylsulfonyl)-2-nitrophenyl]-1,3-benzodioxol-5-amine (MSNBA, GLUT5 inhibitor) or 400 µM phloretin (PT, GLUT2 inhibitor). Basolateral glucose concentration in each experiment was normalized to the value of the NT sample. Basolateral glucose concentration in EM representing the BMI$^{lean}$GA$^{low}$ phenotype was below detection limit. Data from three independent experiments are presented as mean ± SEM.

### Supplementary Materials and Methods

#### Human intestinal enteroid culture establishment and propagation

Small intestinal samples from non-obese healthy (BMI <25 kg/$m^2$), overweight (BMI 25-30 kg/$m^2$) or obese (BMI> 35 kg/$m^2$) patients were obtained during routine endoscopy procedures or from discarded tissue from patients undergoing bariatric surgery (BMI> 35 kg/$m^2$). The BMI classification is based on the World Health Organization guidelines. Based on current National Institutes of Health guidelines, the requirements for BS are as follows: BMI ≥40 kg/$m^2$ or BMI ≥ 35 kg/$m^2$ in patients with obesity-related comorbidities. Detailed information about the patient characteristics can be found at Supplementary Table 1. All methods were carried out in accordance with the approved guidelines and regulations. The experimental protocols were approved by The Baylor College of Medicine institutional review board (protocol numbers H-31793 and H-31910) and Johns Hopkins Institutional review board (protocol number NA_0038329) and the patients' informed consents were obtained. Crypts were isolated from the de-identified tissue samples and enteroid cultures were established as described by our group and others [1, 2, 3, 4]. Briefly, cells were resuspended in Matrigel (Corning) and cultured in a complete medium (CM) with growth factors (CMGF$^+$ medium). CM media contained Advanced Dulbecco's modified Eagle medium/Ham's F-12 (Life Technologies), 100 U penicillin/streptomycin (Life Technologies), 10 mM HEPES (Life Technologies), and 0.2 mM GlutaMAX (Life Technologies). CMFG+ medium is CM medium supplemented with 50% v/v Wnt3A-conditioned medium, 15% v/v R-spondin-1-conditioned medium, 10% v/v Noggin-conditioned medium, 50 ng/ml human epidermal growth factor (EGF) (Life Technologies), 10 nM gastrin I (Sigma), 500 nM A-83-01 (Tocris Bioscience), 10 µM SB202190 (Sigma-Aldrich), 1× B27 supplement (Life Technologies), 1 mM *N*-acetylcysteine (Sigma-Aldrich),10 µM CHIR99021 (Tocris Bioscience) and 10 µM Y-27632 (Tocris Bioscience). CHIR99021 and Y-27632 were removed from CMGF$^+$ media during subsequent media replacements. Conditioned media was obtained from the following cell lines expressing the growth factors: Wnt3A (ATCC CRL-2647 cells (American Tissue Culture Collection), R-spondin-1 (kindly provided by Dr. Calvin Kuo, Palo Alto, CA) and

Noggin[5]. Enteroids were passaged every 10-14 days for expansion or used for monolayer cultures.

**Monolayer plating and differentiation**

Enteroid monolayers were prepared as we previously described in detail [2]. Briefly, transwell filters (24-well nserts, 0.4µM pore polyester membrane, 0.33cm$^2$ surface area) (Corning) were coated with 10µg/cm$^2$ human collagen IV solution at 4ºC overnight. Enteroid fragments were resuspended in CMGF$^+$ media (~50 fragments/100 µl media), and 100 µl of the solution was added to each Transwell filter. CMGF$^+$ media (600 µl) was added to the well of each filter. Confluent monolayers (or 3D cultures in Matrigel) were differentiated for 5-7 days in a differentiation media (CMGF+ media, lacking Wnt3A, R-spondin, and SB202190). Transepithelial electrical resistance (TER) measurements with EVOM2 epithelial ohmmeter (World Precision Instruments) were used to monitor the confluency/differentiation of the monolayers.

**Glucose transport experiments**

Undifferentiated or differentiated confluent enteroid monolayers were washed with mannose buffer (50 mM HEPES-pH 7.4, 138 mM NaCl, 4.7 mM KCl, 1.25 mM $MgSO_4$, 1.25 mM $CaCl_2$, 5 mM Mannose) and incubated at 37 ºC in 5% $CO_2$ for 30 minutes (100 µl added apically to the filter and 600 µl added basolaterally to the well). An aliquot from the basolateral side was taken (as t=0), the same volume of mannose buffer was added, and the apical solution was replaced with 25 mM glucose buffer or 70 mM transport buffer. In glucose buffers, the mannose was replaced with glucose and the osmolarity was adjusted by decreasing NaCl concentration to 128 mM or 105.5 mM. Aliquots were taken at different time points and glucose concentration was determined using the Amplex Red glucose assay kit according to manufacturer's instructions (Thermo Scientific). Phloridzin (Sigma), phloretin (Sigma) or STF-31 (Calbiochem) were used as inhibitors of glucose transporters.

**Gluconeogenesis experiments**

Differentiated confluent enteroid monolayers were washed with 5 mM mannose buffer and incubated for 30 minutes. Apical solution was replaced by 70 mM fructose buffer (5 mM mannose was replaced with 70 mM fructose, and NaCl concentration was decreased to 105.5 mM) or 2 mM Na-pyruvate/20 mM Na-lactate buffer (5 mM mannose was replaced with 2 mM Na-pyruvate and 20 mM Na-lactate, and NaCl concentration was decreased to 107 mM). Cells were incubated at 37 ºC in 5% $CO_2$ for 3 hours and the solutions were collected for glucose measurements.

For 16 hour gluconeogenesis experiments, the solutions were modified based on the Advanced DMEM/Ham's F-12 media formulation and growth factors were included. The differentiated enteroid monolayers were incubated for 30 minutes in 5 mM mannose solution (10 mM HEPES, pH 7.4, 126.6 mM NaCl, 4.15 mM KCl, 0.407 mM $MgSO_4$, 1.05 mM $CaCl_2$, 29.02 mM $NaHCO_3$, 0.3 mM $MgCl_2$, 0.5 mM $Na_2HPO_4$, 0.45 mM $NaH_2PO_4$, 5 mM mannose, 1× B27 supplement, 1 mM *N*-acetylcysteine, 50 ng/ml EGF, 10 nM gastrin I and 500 nM A-83-01). Apical solution was replaced by 70 mM fructose solution (5 mM mannose was replaced with 70 mM fructose, and NaCl concentration was decreased to 94.1 mM). Phloretin (Sigma) or MSNBA (Enamine) were used as inhibitors of fructose transporters.

**Immunofluorescence**

Enteroid monolayers were fixed with 4% paraformaldehyde/PBS for 30 minutes, washed with PBS, permeabilized and blocked in PBS containing 15% FBS, 2% BSA, 0.1% saponin for 45 minutes. The cells were immunostained using the following antibodies (1:100 dilution) at 4ºC overnight: Glut1 (ab15309, Abcam), SGLT1 (07-1417, Millipore), and occludin (331500, Life Technologies). Cells were washed with 1x PBS and incubated with the following secondary antibodies (Life Technologies) at room temperature for 1 hour: Alexa Fluor 488 donkey anti rabbit, Alexa Fluor 555 donkey anti mouse, Wheat germ agglutinin-555 (WGA), Phalloidin-633 or Hoechst. Cells were washed with PBS, filters were cut and mounted on glass slides using Fluor-save reagent (Millipore). Fluorescent confocal imaging was performed using Zeiss 510 LSM (Zeiss) and the images were analyzed using Metamorph software (Molecular Devices).

**Immunoblot analysis**

Cells from either 3D Matrigel cultures or monolayers were harvested, resuspended in lysis buffer (60 mM HEPES pH 7.4, 150 mM KCl, 5 mM Na3EDTA, 5 mM EGTA, 1 mM Na3VO4, 50 mM NaF, 1% Triton X-100) with protease inhibitor cocktail (Sigma-Aldrich) (1:100) and disrupted by sonication. Protein concentrations were determined using the BCA Protein Assay (Thermo Scientific). Total cell lysate (25 µg) was mixed with Laemlli loading buffer containing SDS/β-mercaptoethanol and incubated for 10 minutes at room temperature. Proteins were separated using Tris-Glycine gels (Thermo Scientific) and wet-transferred to a nitrocellulose membrane at 100 V for 40 minutes using transfer buffer (34 mM Tris, 39 mM glycine, 0.004% SDS, 20% methanol). Membranes were blocked using 5% milk/Tris-buffered saline with 0.1% Tween (TBST) for 1 hour. Membranes were washed with 1x TBST and incubated overnight at 4 ºC with the following primary antibodies diluted in 3% BSA/TBST: GLUT1 (1:400, ab15309-Abcam), PEPCK1 (1:400, HPA006507-Sigma), GAPDH (1:1000, G8795-Sigma). Membranes were washed with 1x TBST and incubated with IRdye-conjugated secondary antibodies diluted 1:10.000 in 1x TBST. Membranes were imaged using an infrared imaging scanner (Li-Cor). The expression levels of the target proteins were obtained by normalizing the relative fluorescence intensity of the target protein to the fluorescence intensity of GAPDH.

SGLT1 blotting was performed using a modified protocol with cell membrane lysates[6, 7]. Enteroids were harvested and resuspended in 25 mM HEPES, pH 7.4; 250 mM sucrose; 1 mM EDTA solution with protease inhibitor tablets (Roche). Cells were disrupted by using a 1 ml syringe with 25 gauge hypodermic needle. The cells were passed through the needle 10 times, and the lysate was centrifuged at 500 x g for 10 min to remove cell debris and nuclei. The supernatant was again centrifuged at 3,000 x g for 10 min to remove mitochondria. The resulting supernatant was centrifuged at 150,000 x g for 90 minutes to pellet the total cell membranes which were resuspended in 150 mM mannitol, 2.5 mM EGTA, 12 mM Tris/HCl, pH 7.4. All the centrifugation steps were performed at 4°C. Total cell membrane (50 µg) was mixed with non-reducing Laemlli loading buffer, denatured for 15 min at 65 °C and proteins were separated using 4-12% NuPage gels (Thermo Scientific) using 1x MOPS buffer. Proteins were wet-transferred to an Immobilon PVDF membrane (Millipore), and membranes were blocked in blotting buffer (5 % nonfat dry milk, 0.15 M NaCl, 1% Triton X-100, and 20 mM Tris·HCl, pH 7.4) for 1 hour. Na/K-ATPase α1-subunit (1:1000, 05369-Millipore) and SGLT1 antibody (1:500, kindly provided by Dr. Hermann Koepsell, University of Würzburg, Germany) were diluted in the blotting buffer and membranes were incubated at 4°C overnight. Washing was performed using blotting buffer, and fluorescence dye-conjugated secondary antibodies diluted 1:10.000 in blotting buffer were added. Membranes were washed with blotting buffer, followed by a wash using 20 mM Tris·HCl buffer, pH 7.4 and imaged using the Odyssey Infrared Imaging Scanner (Li-Cor)

**Real time polymerase chain reaction**

Total RNA was extracted from 3D Matrigel or monolayer cultures using the PureLink RNA Mini Kit (Ambion) according to the manufacturer's protocol. Complementary DNA (cDNA) was synthesized using SuperScript VILO MasterMix (Invitrogen). Quantitative PCR was performed on a QuantStudio 12K Flex Real-Time PCR System (Applied Biosystems). Expression of GLUT2, GLUT5, and glucose-6-phosphatase was determined with 18S rRNA as endogenous control. Each sample was run in triplicates. The primer sequences were as following: GLUT2-Fwd (TCTCTGCTACTCTTTTTCTGTCCA), GLUT2-Rev (TCATATCCTCTGA GTCTTTTCAAGC), GLUT5-Fwd (TCTCCTTGCAAACGTAGATGG), GLUT5-Rev (GAAGA AGGGCAGCAGAAGG), Glucose-6-phosphatase-Fwd (AAGCCGACCTACAGATTTCG), Glucose-6-phosphatase-Rev (CGTGACAGACAGACATTCAGC), 18S-Fwd (GCAATTATTC CCCATGAACG), 18S-Rev (GGGACTTAATCAACGCAAGC).

**Lucifer Yellow Paracellular Permeability**

Differentiated confluent enteroid monolayers were washed with 5 mM mannose buffer and incubated for 30 minutes. Apical solution was replaced by 5 mM mannose buffer containing 200 µM lucifer yellow. Cells were incubated at 37 ºC in 5% $CO_2$ for 2 hours and the basolateral solutions were collected for fluorescence measurements.

### Statistical analysis

Statistical analyses were performed using Prism (Graphpad). Unpaired Student's t-test was used to analyze for statistical differences with $p<0.05$ considered statistically significant. The data are expressed as mean ± S.E.M, unless otherwise stated.

### Acknowledgments

This work was supported by NIH grants P01 AI125181. We also acknowledge the Integrated Physiology and Imaging Cores of the Hopkins Conte Digestive Disease Basic and Translational Research Core Center (P30 DK089502). The authors acknowledge the Kudsi Imaging Facility and the assistance of John Gibas as well as the Integrated Physiology and Translational Research Enhancement Cores of the Hopkins Conte Digestive Disease Basic and Translational Research Core Center. The authors thank James Potter and Dr. Steven Brant (Hopkins) and Xi-Lei Zeng (Baylor) for their assistance in acquiring patient specimens from which proximal small intestinal enteroid lines were established. We thank Dr. Mark Donowitz for his helpful suggestions and critical reading of the manuscript.

### Contributions

N.M.H., J.Y., K.F.J., and N.W.B. performed the experiments. N.M.H., N.C.Z. and O.K. designed the experiments. N.M.H., N.C.Z., and O.K. analyzed the results. V.S. and V.K. acquired the patient specimens. N.M.H. and O.K. prepared the manuscript. N.M.H., S.E.B., M.K.E., N.C.Z., and O.K. discussed the results and edited the manuscript. The final manuscript was approved by all authors.
